# Cross-species modeling and enhancement of cognitive control with striatal brain stimulation

**DOI:** 10.1101/2024.02.16.580680

**Authors:** Adriano E Reimer, Evan M Dastin-van Rijn, Jaejoong Kim, Megan E Mensinger, Elizabeth M Sachse, Aaron Wald, Eric Hoskins, Kartikeya Singh, Abigail Alpers, Dawson Cooper, Meng-Chen Lo, Amanda Ribeiro de Oliveira, Gregory Simandl, Nathaniel Stephenson, Alik S Widge

## Abstract

Brain disorders, particularly mental disorders, might be effectively treated by direct electrical brain stimulation, but clinical progress requires understanding of therapeutic mechanisms. Animal models have not helped, because there are no direct animal models of mental illness. We show a path past this roadblock, by leveraging a common ingredient of most mental disorders: impaired cognitive control. We previously showed that deep brain stimulation (DBS) improves cognitive control in humans. We now reverse translate that result, showing that DBS-like stimulation of the mid-striatum improves cognitive control in rats. Using this model, we identify a mechanism, improvement in domain-general cognitive control, and rule out competing hypotheses such as impulsivity. The rat findings explain prior human results and have immediate implications for clinical practice and future trial design.

**One Sentence Summary:** Developing a reliable animal model of a human brain stimulation therapy reveals that this therapy works by enhancing the brain’s ability to process conflicting pieces of evidence.

## Introduction

Deep brain stimulation (DBS) uses surgically implanted electrodes to modulate target brain circuits. DBS is highly successful in movement disorders (*1*) and is being actively investigated for treatment of psychiatric disorders, including treatment-refractory obsessive-compulsive disorder (OCD) (*2*), Tourette syndrome (*3*), and major depressive disorder (MDD) (*4*). In all these disorders, DBS’ mechanism of action remains unclear (*1*, *5*, *6*). The lack of mechanistic understanding leads to difficulties in programming stimulation to individual patients’ needs, which contributes to clinical trial failures (*5*, *6*). Human neurophysiology studies offer hope for better stimulator programming (*6–9*) but are similarly limited by the internal heterogeneity of psychiatric diagnoses (*6*, *10*, *11*). In movement disorders, mechanistic understanding arose from animal models (*12*), and those models are testbeds for new treatment paradigms (*13*, *14*). In contrast, psychiatric syndromes are difficult to model in animals, limiting therapy development (*15*, *16*).

Diagnostic heterogeneity might be overcome, and reliable animal models created, by modeling constructs that cut across and form the basic ingredients of psychiatric illnesses (*10*, *16*, *17*). Cognitive control is a particularly promising construct for modeling DBS, because (1) it is highly relevant to multiple illnesses and (2) it can be modified via DBS. Cognitive control is the ability to adjust and regulate thoughts and behavior in service of a specific goal (*18*, *19*). It involves the ability to suppress or override prepotent responses in favor of more adaptive responses. Cognitive control depends on a distributed brain network connecting prefrontal regions with the basal ganglia, striatum, and thalamus (*20–22*). Impaired cognitive control is linked to a wide range of psychiatric disorders, including MDD (*23*), OCD (*24*, *25*), and addictions (*26*).

Multiple studies have enhanced cognitive control with DBS-like stimulation of the human striatum and internal capsule, with correlation between that enhancement and improvement in mood/anxiety symptoms (*27–29*). Those studies were limited by windows of opportunity in clinical volunteers. They did not verify that the effects were repeatable or demonstrate mechanisms. The last point is critical, because the claimed cognitive control enhancement was a decrease in response times (RTs) on a psychophysical task. An RT decrease might reflect improved cognitive control, but might also represent a deleterious effect, such as impulsivity. If the task was sufficiently easy, participants might have been able to increase their speed without increased caution.

Conversely, cognitive control has multiple subcomponents (*19*, *30*, *31*). The RT effect might reflect improvement in motivation, attentional control to focus on task demands, pre-commitment to high-control behaviors (proactive allocation), adjustment to changing requirements (reactive allocation), or conflict processing (domain-general control). All are consistent with the subjective effects of striatum/capsule stimulation, including increased: motivation (*32*, *33*), ability to refocus attention away from distress (*27*, *34*), ability to effortfully resist symptoms (*35*), and drive towards pleasure and social interaction (*36*, *37*). Recipients also report less concern for negative outcomes (*34*, *37*). Animal models could adjudicate between those mechanisms. Rodent models would be particularly useful, as rodents have a wide range of toolkits for manipulating specific neuronal subpopulations and circuits (*38*, *39*). Further, there is rodent-human homology in circuits believed to underpin cognitive control (*40*, *41*). Related executive functions such as extinction learning are improved by stimulation of these homologous structures (*42*, *43*).

We leverage that homology to develop a reverse translational rodent model of psychiatric DBS that reveals underlying mechanisms. First, we show that human RT enhancements can be replicated in rodents by stimulation of the mid-striatum and embedded corticofugal fibers. This enhancement replicates a dorso-ventral anatomic gradient observed in humans. We then apply this model to dissect DBS’ cognitive effects. We explore the potential mechanisms above, and through multiple behavioral paradigms and computational modeling, show that DBS improves task performance by increasing the efficiency of conflict adjudication (domain-general cognitive control). Physiologically, we link this improvement to the simultaneous modulation of multiple prefrontal regions, possibly via their cortico-thalamic projections. Closing the translational circle, we demonstrate that the same cognitive mechanism is present in the original human data, i.e. the animal model has explanatory power for clinically relevant phenomena.

## Results

### Replicating DBS’ effects on cognitive control in rodents

We probed cognitive control in rats using a Set-Shift task where rats shift between cue (Light) and spatial (Side) rules (Figure 1A-D, Table S1) (*44*, *45*). Human cognitive control enhancement depended on stimulating specific sites in the mid/dorsal striatum and internal capsule (*27*). We mapped the effects of DBS-like stimulation at homologous sub-regions of rat striatum (n=35, Figure 1E, Figure S1). This model replicated the human findings. Mid-striatum stimulation significantly improved RTs (β = -34 ms, p < 2.7e-17); other sites did not (p > 0.05, Figure 1F, Table S2). As in humans, there was no significant effect on response accuracy or any other outcome (p > 0.05, Figure 1G, Figure S2C-D, Tables S3-S5). The improvement was consistently present across animals and days (Figure S2E-F). It was not explained by changes in general activity (p > 0.05, Figure S3A-C, Table S6-S10), motivation (p > 0.05, Figure S3D-E, Tables S9-S10) or lesion effects (p > 0.05, Table S2). It required the higher frequencies generally used for DBS. 20 Hz stimulation worsened the total errors (β=0.23, p=5.50e-4, Figure S4, Table S11) and RTs (β = 25 ms, p=2.96e-4, Figure S4, Table S12), in an independent cohort that also replicated the results of the initial 130 Hz study (β = -48 ms, p=1.54e-34, Figure S4, Table S12).

**Figure 1.**
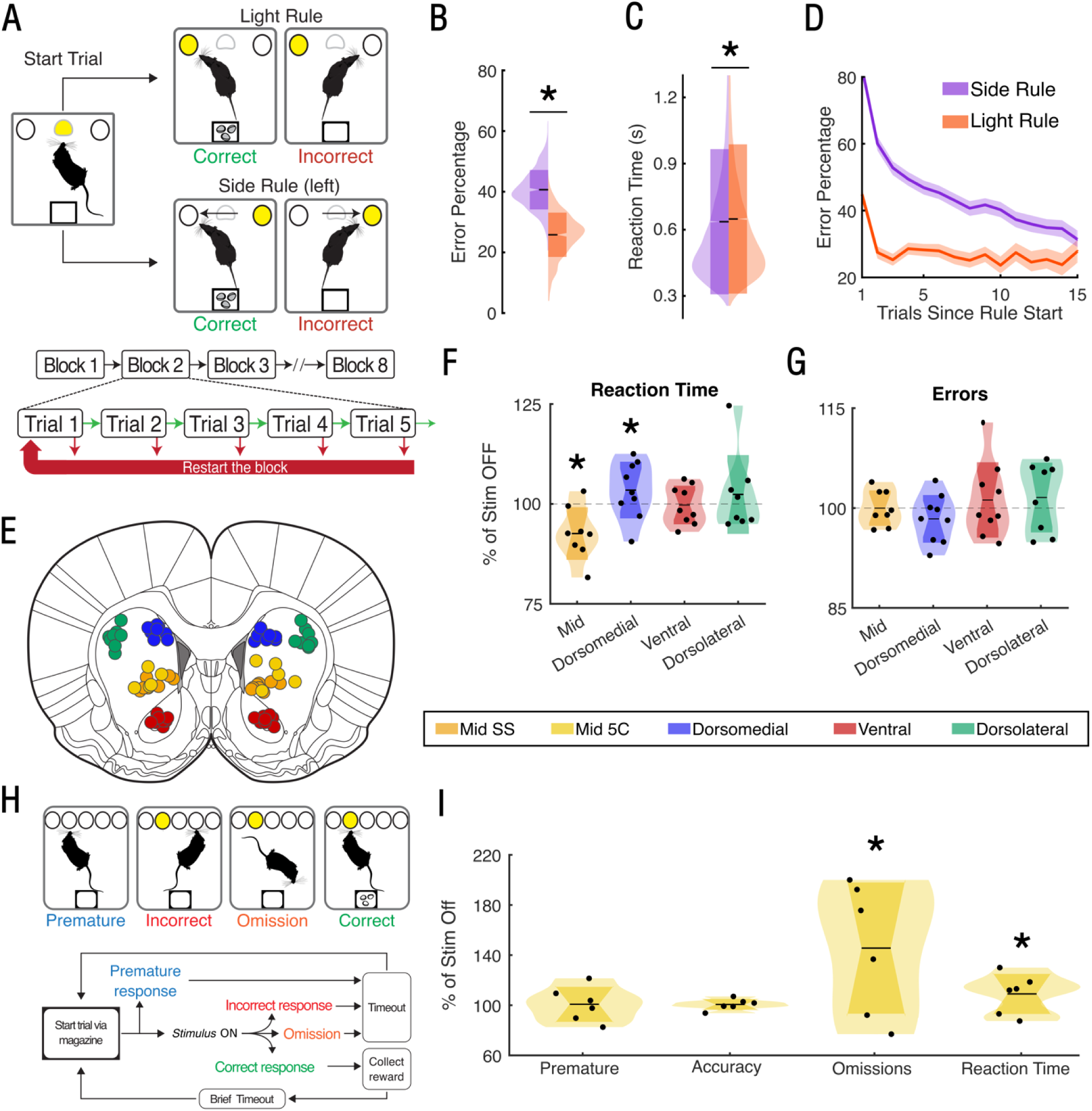
Behavioral testing paradigms and stimulation outcomes. (A-D) Set-Shift task. (A), task schematic. Rats must nosepoke to either an illuminated hole (Light rule) or ignore the light and always nosepoke on a specific side of the chamber (Side rule). The rules are not cued, and shift after the rat has completed 5 correct trials of the current rule. Errors reset to the beginning of the current 5-trial block. (B), probability of error based on the current rule. The cue light acts as a pre-potent stimulus that is difficult to ignore, leading to increased errors during Side blocks (β=0.64, p<0.001). (C), RT as a function of rule. Because Side rules do not require sensory processing (the correct answer is known even before the cue light is seen), they have slightly faster RTs (β=12 ms, p<0.001). (D), probability of error after a rule shift. In both rules, rats learn the task and progress through blocks, evidenced by decreasing error rate as a block proceeds. However, as in (B), Side rules are more difficult and the error does not rapidly asymptote, whereas Light rules can be reliably learned in 2-3 trials. (E-G), stimulation experiment. (E), histologically confirmed sites of implant for sub-regions of the striatum. The Mid-striatal target was the only target tested during the 5-choice experiment (see below), whereas all 4 targets were tested for the primary Set-Shift experiment. (F), distributions of RT as a function of stimulation type, expressed as a percentage of stimulation-off. Distributions are computed over rats; each dot shows the median for one rat. Mid-striatal stimulation specifically replicates the effect seen in prior human data stimulating at this target (see main text; β= -34 ms, p < 0.001), whereas dorsomedial striatal stimulation worsens RT. (G), distributions of error count (during the whole session) as a function of stimulation type, showing no change, which also replicates prior human results. (H-I), 5-choice serial reaction time task (5-CSRTT). (H), task schematic. A trial began when a rat entered the food magazine. After a 5-second interval, a brief light stimulus was presented, pseudorandomly, in one of five different nosepoke ports on the opposite wall. If the rat poked the lit aperture, it received a food pellet followed by a 5-second timeout (“correct response”). If the rat responded before the light was presented (“premature response”), in an incorrect aperture (“incorrect response”) or did not respond within the defined time period (“omission”) the house light extinguished briefly to signify failure, the rat received no reward and a 5-second timeout. (I), outcomes, following the plotting conventions of (F-G). Mid-striatal stimulation did not increase premature responses or decrease accuracy, i.e. it did not show an impulsivity effect. It did significantly increase omissions (β = 0.448, p < 0.001) and increase RT (β = 49 ms, p < 0.001), both of which are opposite to what would be expected from impulsive behavior. All p-values represent Wald Z-tests of model parameters from generalized linear mixed models (GLMMs), see Methods for formulae and Tables S1-S3 (Set-Shift) and S17-S20 (5CSRTT) for statistical details.

### RT improvements are not impulsivity

We tested for increased impulsivity or motivation using the 5-Choice Serial Reaction Time Task (5CSRTT, n=6, Figure 1E/H, Figure S5, Tables S15-S16) (*46*). There was no change in premature responses, the most common impulsivity metric (p=0.35, Figure 1I, Table S17). 5CSRTT RTs <u>increased</u> (β = 49 ms, p=0.005, Figure 1I, Table S18), which may reflect improved cognitive control in a situation where response withholding is essential. Stimulation also increased omissions (β = 0.45, p=1.06e-8, Figure 1I, Table S19), arguing against an increase in general motivation.

### Mid-striatal stimulation non-specifically increased proactive control

Cognitive control involves both planned, proactive adaptive responses and reactive cancellation of maladaptive responses (*19*, *31*). We examined whether mid-striatal stimulation affected either process. Rats frequently nose-poked during the inter-trial interval (ITI), and we reasoned that these might reflect rehearsal or planning (proactive control). Supporting that concept, during the ITI rats showed both early nosepokes that were more common after incorrect responses, then a ramping pattern of late-ITI nosepokes that were more common after correct responses (Figure 2A-B). After incorrect responses, ITI nosepokes suggested correction/regret – they were commonly directed to the port that would have been correct (β = 0.53, p = 7.09e-18, Figure 2A, left inset, Table S21-S22). Similarly, after correct responses, nosepokes were more common in the port that was just rewarded (β = 1.89, p = 7.98e-302, Figure 2B, right inset, Table S23). Both could reflect proactive control, because on Side blocks, it is possible to plan the correct response before a trial begins. Thus, ITI pokes might reflect rehearsal to keep a desired response in working memory. Consistent with this thesis, rats were more likely to poke a side they had rehearsed during the immediately preceding ITI – but only if they ignored the cue light and chose the non-illuminated port (β = 1.84, p=4.50e-129, Figure 2C, Table S24). Further, late-ITI nosepokes in the middle (trial initiation) port predicted faster RTs on the subsequent trial (β = -36 ms, p=1.66e-15, Figure 2D, Table S25). Also consistent with a proactive control effect, this RT improvement was augmented by rehearsal. If a rat poked both the middle and the port it would choose on the upcoming trial (a “Choice” poke), the RT improvement was greater (β = -12 ms, p=0.021, Figure 2D, Table S25), whereas nosepokes into a port discordant with the upcoming trial behavior (“Non-Choice” pokes) had no effect beyond that of the accompanying middle-port poke (p=0.4448, Figure 2D, Table S25). Proactive rehearsal did not, however, explain the RT effect of stimulation, because stimulation did not specifically increase it. Rather, mid-striatal stimulation increased all forms of ITI pokes (all p < 6.42e-3, Figure 2E, Table S26), including Non-Choice pokes unrelated to cognitive control.

**Figure 2.**
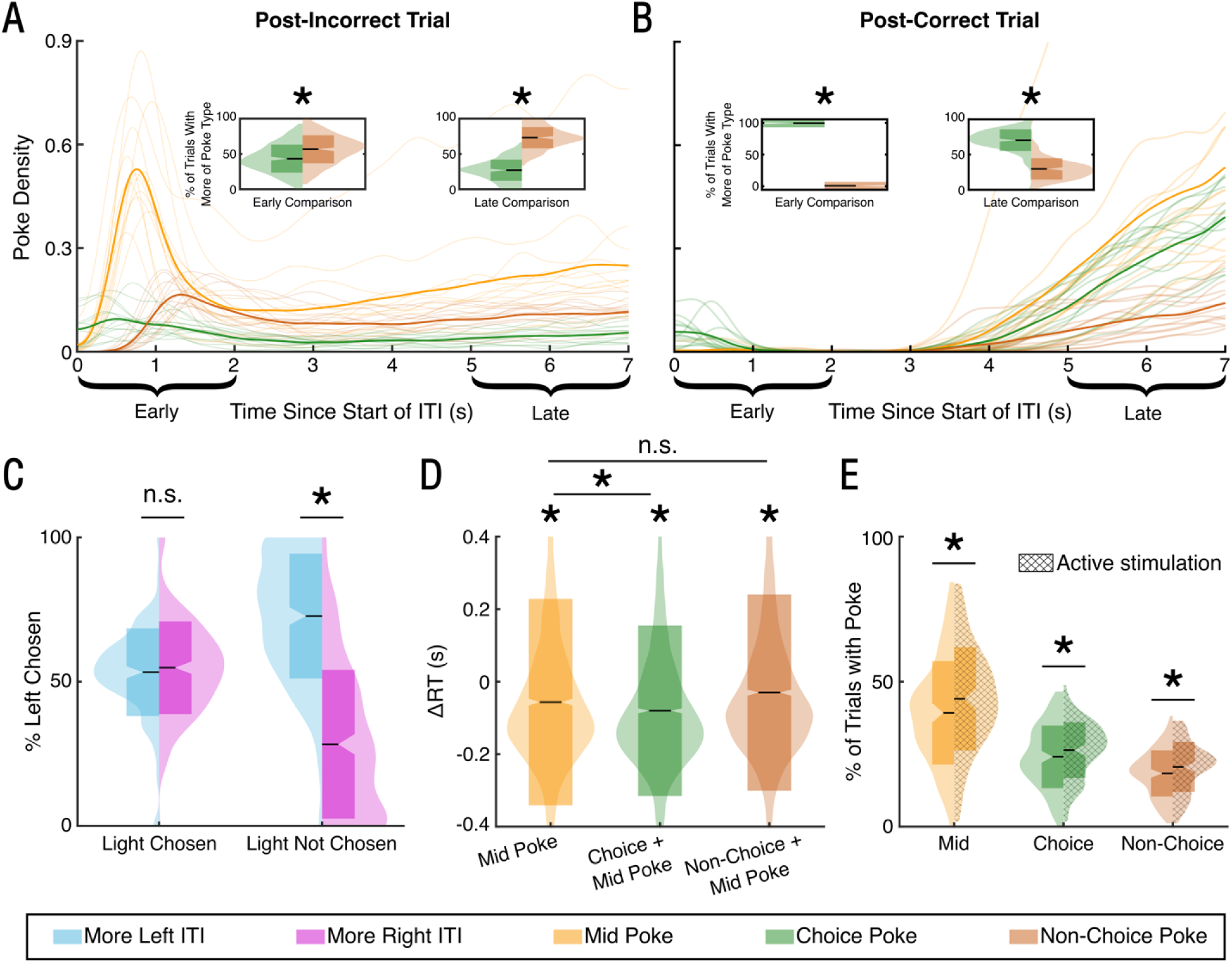
Precommitment related behaviors during inter-trial intervals (ITIs). (A-B) probability of poking the mid (trial initiation) and side ports following a decision. Thick lines show the mean over all animals and sessions, while light shaded lines in background show individual animals. Each plot is a kernel density estimate of the distribution, using a Gaussian kernel with 0.2s standard deviation. “Choice” and “Non-choice” pokes are pokes in the ports that were and were not chosen in the preceding trial, respectively. We specifically emphasize “Early” (first 2 seconds) and “Late” (last 2 seconds) behavior. Following an incorrect response (A), rats showed an Early ITI peak in poking that leveled to a stable rate for the remainder of the inter-trial interval. Pokes in the side ports were significantly more likely to match the option that was not chosen in the preceding trial, both in the Early (left inset (A), β=0.53, p<0.001, Table S22) and Late ITI (right inset (A), β=1.73, p<0.001, Table S22). These resemble regretful correction and rehearsal, respectively. Following a correct response (B), rats made fewer Early pokes (β =-0.24, p<0.001, Table S21), but increased their poking in the Late period (β =0.12, p<0.001, Table S21). Pokes in the side ports were significantly more likely to match the option that was rewarded in the preceding trial both in the Early ITI (left inset (B), β=8.27, p<0.001, Table S23) and the Late ITI (right inset (B), β=1.89, p<0.001, Table S23). These are consistent with rehearsal of a rewarded option. (C-D) Effects of ITI behavior on the subsequent trial. In these panels, “Choice” and “Non-Choice” now refer to the upcoming trial, rather than the preceding trial as in (A-B). (C) ITI pokes affected subsequent trial behavior (precommitment), but only when the rat chose to ignore the cue light (e.g., following successful learning of a Side rule). Distributions show the probability of choosing the left or right port, based on which port was most often poked during the Late portion of the preceding ITI. If the lit port was chosen, ITI pokes had no effect (β=-0.087,p=0.08, Table S24). If the lit port was not chosen, rats were much more likely to poke the port that they had rehearsed during the ITI (β=1.84,p<0.001, Table S24). (D) ITI pokes in the late portion of the ITI affected subsequent decision times, also consistent with precommitment. Plots show the distribution of RTs conditioned on different types of ITI behavior. Pokes in the mid port significantly lowered RTs (yellow, β1=-0.04,p<0.001, Table S25), but there was an even stronger effect (green, β2=-0.01,p=0.02, Table S25) when the rat had rehearsed a full response sequence, namely a mid poke along with a poke into the side port that matched the subsequent choice. In contrast, there was no additive effect of a mid poke along with a poke into the side that was subsequently not chosen (orange, β3=0.00, p=0.44, Table S25). * above individual distributions indicates a mean different from the distribution when no ITI pokes occurred. (E) Distributions of late ITI poking into the mid and side ports with (hatched) and without mid-striatal stimulation. Stimulation increased all poke types, both those consistent with precommitment (choice pokes) (green, β=0.21, p<0.001, Table S26) and those that might represent general motivated behavior (mid (yellow, β=0.24,p<0.001, Table S26) and non-choice pokes (orange, β=0.09, p<0.007, Table S26)). * above each distribution reflects a significant difference between stimulation on and off. All p-values represent Wald Z-tests of model parameters from generalized linear mixed models (GLMMs), see Methods for formulae.

### Stimulation improves cognitive control by improving decisional efficiency

The RT decrease could also reflect reactive adaptation or a general improvement in conflict resolution/evidence. To explore these, we fit a hybrid model (*47*) to the behavior data (Figure 3A, Figure S6), incorporating both reactive reinforcement learning (RL, (*31*, *48*)) and diffusion-based evidence accumulation (DDM, a model of conflict resolution (*49*)). The RL component included an adaptive learning rate and forgetting function, based on a model selection analysis (Figure S7A). Simulated responses from the RLDDM strongly matched empirical behavior and psychometrics (Figure 3B-D, S7B-E). The ITI behaviors in Figure 2 were captured in the RLDDM’s bias term, which incorporates value information specifically related to the Side rule (Figure 3D). Mid-striatal stimulation did not alter any learning components of the model, arguing against a reactive control mechanism (Figure 3E). Rather, it altered evidence accumulation components in a specific fashion that promotes efficient conflict resolution. Stimulation lowered the boundary separation term (pd=99.85%, Std.Median=-0.3335, 1.05% in ROPE), which reflects the amount of evidence needed to reach a choice. This on its own would decrease RTs but also increase errors, because narrower boundaries are more easily reached by noise. However, stimulation also increased the drift rate (pd=99.25%, Std.Median=-0.4406, 2.925% in ROPE), which drives the model towards the correct-choice boundary. When combined with tighter bounds, this leads to more efficient conflict resolution, where the system needs less time to reliably reach the correct choice. There was a much smaller stimulation effect on the bias term (pd=88.525%, Std.Median=0.296, 15.425% in ROPE), which connects the RL component to the DDM. These three factors explained nearly 100% of the stimulation effect (Figure 3F). This modeling implicates the general process of conflict resolution, rather than specific strategies for cognitive control allocation, as the primary mechanism of cognitive control enhancement.

**Figure 3.**
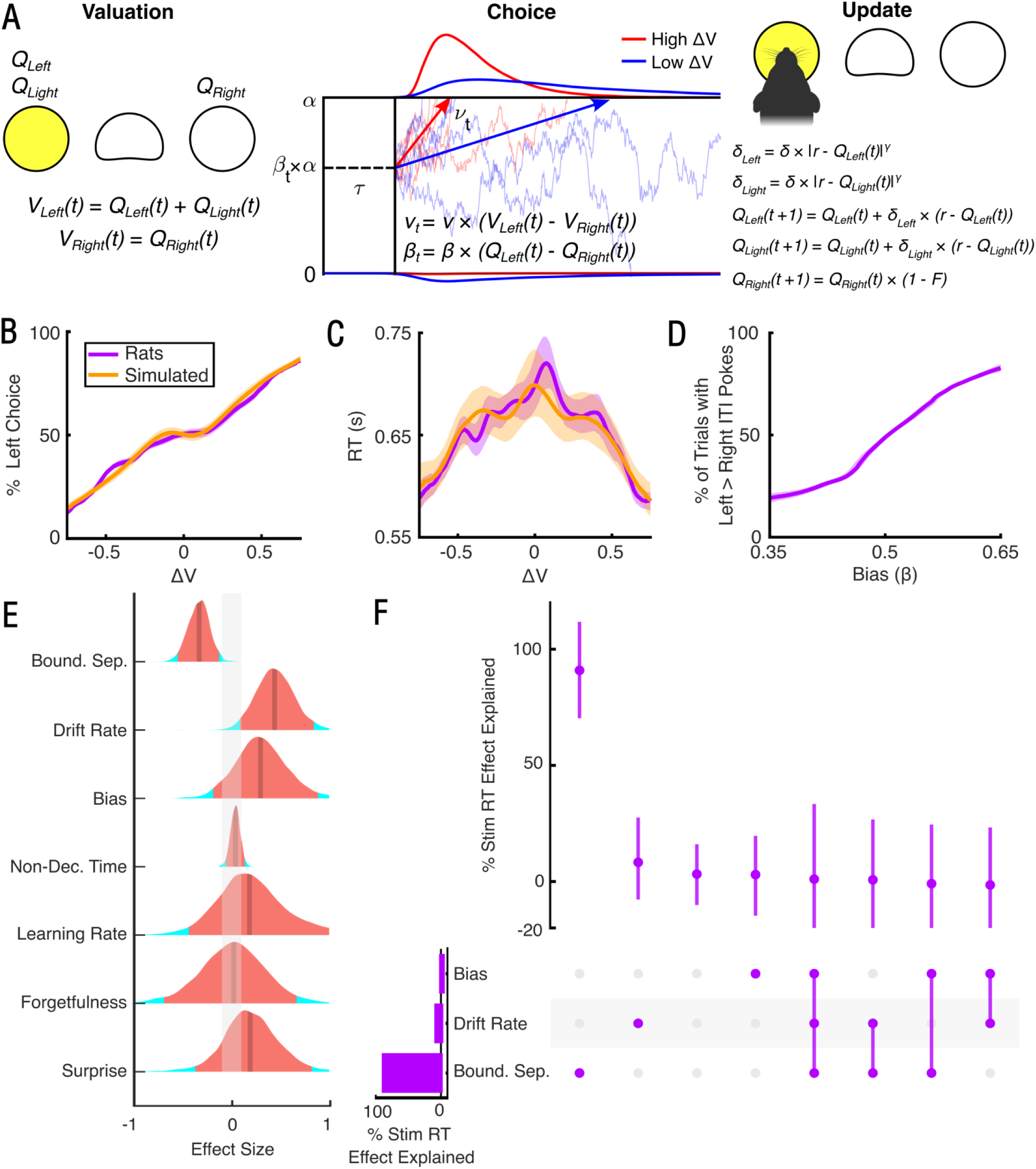
Mid-striatal stimulation decreases reaction times specifically by improving evidence processing and conflict resolution. (A) Illustration of the Reinforcement Learning-Drift Diffusion Model (RLDDM) fit to Set-Shift behavior. The RLDDM first assigns values V to the ports based on its underlying estimates Q of the current value of the sides (left, right) and the light. The model then chooses a port based on a drift-diffusion process described by non-decision time τ, drift rate ν, and decision boundary α. The difference in total value informs the drift rate, while the difference between the sides alone informs the bias ß. Trials with high value difference are likely to have shorter reaction times and more consistently correct choices (red vs. blue curves above). After the port choice, values Q are updated based on a reinforcement learning process, with a non-linear scaling factor γ to diminish small errors and accentuate large ones. The value of the unchosen option decays according to a forgetfulness factor F. The model was fit hierarchically to the data from all rats, allowing individual-specific estimates of model parameters and their change with stimulation. (B-D) Posterior predictive simulations of behavioral outcomes for models (orange) and rat behavior (purple) over 4000 hierarchical posterior draws; see Methods. Shading shows the 95% highest density interval across animals and trials. (B) Choice as a function of the difference in value between the left and right sides (ΔV), where higher X-axis values represent a higher value for the left port. Consistent with expected psychometrics, rats and model simulations were both substantially more likely to select the more valuable side, with a small plateau of equiprobability around the ΔV=0 equipoise point. (C) RT as a function of the difference in value between the left and right sides, with same conventions as (B). Reaction times were greater for smaller value differences than larger ones, for both rats and simulated model data. This is consistent with greater decision difficulty as the choices become near-equivalent. (D) Probability of an ITI poke as a function of the model’s bias term. Higher X-axis values correspond to greater bias ß towards the left port, while the Y-axis shows the percentage of left vs. right pokes during the ITI. As the bias term increased, so did rats’ tendency to rehearse the left port during the ITI. This bias term can therefore be treated as a smoothed estimate of the planning/rehearsal behaviors during ITIs (Figure 2). The simulated and empirical curves entirely overlap. (E) Distributions of stimulation effect on model parameters over 4000 posterior draws. The median of the distribution is indicated by the solid line while the 95% highest density interval is in orange. The grey shaded area around zero represents the Region of Practical Equivalence (ROPE) for a null effect (effect size less than 0.1). Electrical stimulation of mid-striatum substantially decreased boundary separation α (d=-0.33, pd=0.999, 0% in ROPE), substantially increased drift rate ν (d=0.44, pd=0.993, 0.6% in ROPE), and marginally increased bias ß (d=0.30, pd=0.885, 15.4% in ROPE). None of the other parameters showed meaningful effects with stimulation. As described in the main text, the interaction of these parameter changes represents an increase in general cognitive control. (F) UpSet plot illustrating the percentage of the stimulation-related decrease in RT explained by stimulation effects on RLDDM model parameters. Each row corresponds to a parameter, with the bar indicating the total percentage of the stimulation-RT effect explained by that parameter. Each column corresponds to an intersection between parameters with the filled in dots indicating which parameters are part of the intersection. The upper panel indicates the percentage of the stimulation-RT effect that is unique to that intersection. The median and 95% highest density interval across posterior draws from the model are shown by the point and the bar respectively. The stimulation related decrease in reaction time was fully explained by unique contributions from boundary separation (91%, HDI=[70,112]) and drift rate (10%, HDI=[-8,28]) with negligible effects from the interaction between these or other parameters.

### Stimulation effects on cognitive control correlate with modulation of prefrontal cortex

Prominent theories argue that stimulation of the capsule/striatum works primarily by retrograde modulation of prefrontal cortex (PFC) (*50–52*), with mixed excitatory-inhibitory effects (*28*, *53*). Consistent with this model, striatal stimulation increased c-fos expression throughout PFC. (Figure S8A-B). The improvements modeled in Figure 3 appeared to arise from modulation of prelimbic (PL) and infralimbic (IL) cortices, as in these regions c-fos expression correlated moderately (r >0.24) with RLDDM parameters. Boundary and drift rate changes were separable, with the former loaded more onto PL and the latter more onto IL modulation (Figure S8C).

### Decisional efficacy changes translate across species

We finally tested whether these rodent findings explained the original human cognitive control improvements (*27*, *28*). In those studies, participants performed a Multi-Source Interference Task (MSIT, Figure 4A), where trials switched rapidly and unpredictably between two types. We thus fit DDMs without the RL component (Figure 4B) to the participants from (*27*) who received the mid/dorsal striatal stimulation that most improved RT (Figure S9). Stimulation again increased the drift rate (Figure 4C) (pd=0.978, Std.Median=1.31, 1.47% in ROPE), and this effect was present in 4/5 participants. Model selection argued against a change in the boundary separation (Figure S9B), as would be expected given the differing structure of the rodent and human tasks (see Methods, Discussion). We replicated these findings in an independent dataset from (*28*) (Figure S10; pd=0.996, Std.Median=0.62, 1.03% in ROPE).

**Figure 4.**
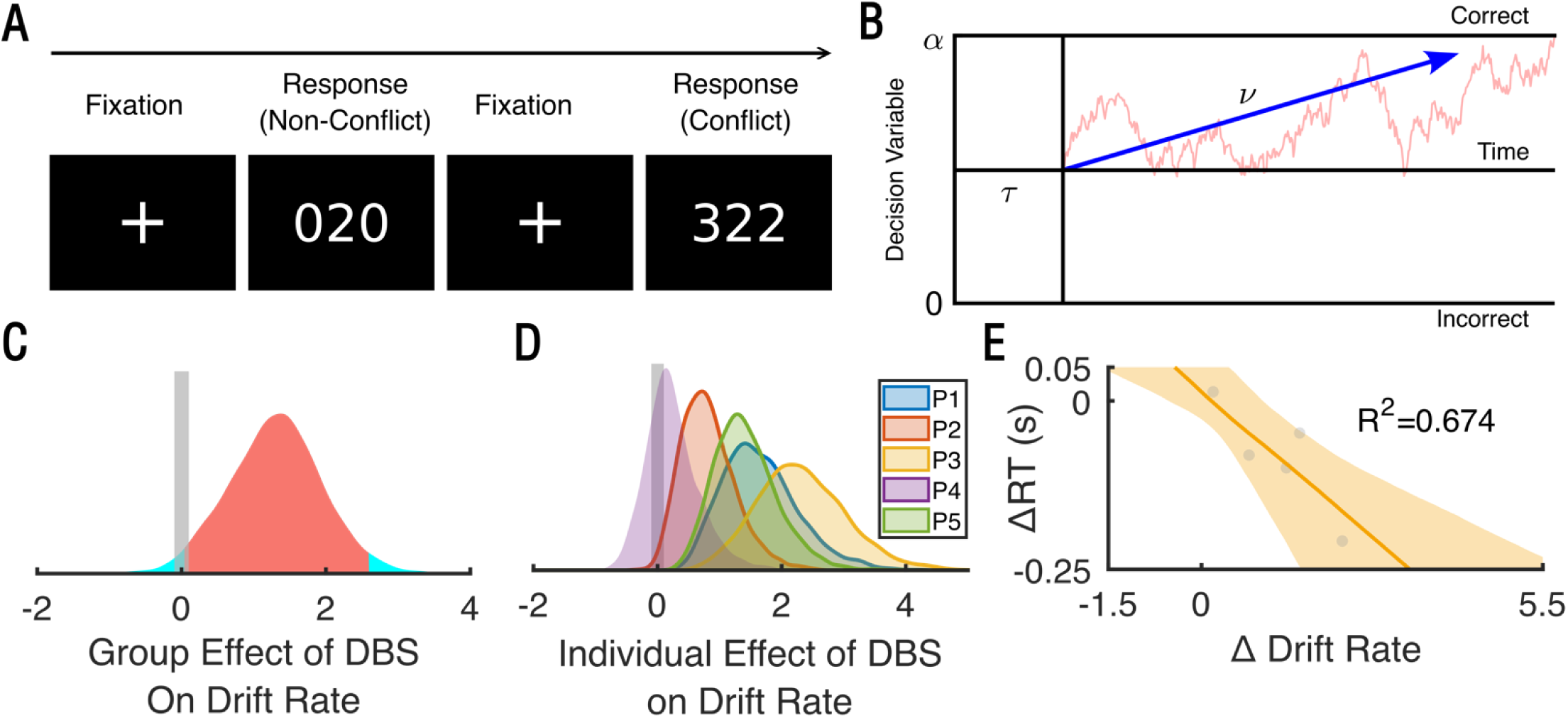
Changes in evidence processing identified in rats also explain the previously reported human effects. (A), schematic of the Multi Source Interference Task (MSIT). Participants choose the number that is different from the other two. On Conflict trials, the target number is out of position and flanked by distractors that are valid targets, leading to interference that requires cognitive control to suppress (*27*, *65*). In Non-Conflict trials, the target is in position and flankers are not targets. In (*27*, *28*), we reported that striatal/internal capsule stimulation improves RT on this task. (B), model schematic. We fit a standard drift-diffusion model (DDM), exploring the potential effect of striatal stimulation on boundary separation α and/or drift rate ν. Based on Figure 3’s results, we did not model potential effects on non-decision time τ. (C) Distributions of stimulation effect on drift rate over 4000 posterior draws. The median of the distribution is indicated by the solid line while the 95% highest density interval is in orange. The grey shaded area around zero represents the Region of Practical Equivalence (ROPE) for a null effect (effect size less than 0.1). Electrical stimulation of mid-striatum substantially increased the drift rate (d=1.31 pd=0.978, 1.47% in ROPE). (D), individual distributions of drift rate changes, over 4000 posterior draws, on the same scale as (C). In 4 of the 5 participants, stimulation substantially increased drift rate. (E), correlation between drift rate change and stimulation-induced RT change. Each dot represents one participant, specifically the RT change plotted against the point estimate of the drift rate change (median of the distributions shown in panel D). The solid line shows a line of best fit. The shaded region represents the 95% highest density interval of this line. We calculated this by running a separate correlation for each posterior draw (distributions in panel D) against the change in median RT.

## Discussion

We developed a reverse translational model of psychiatric DBS, with homology to and explanatory power for effects previously shown in humans. This model highlights the power of a cross-diagnostic/domain-focused approach. There are no strong animal models of psychiatric illness, in part because the core features of those illnesses are subjective self-reports (*15*, *16*). Experts have repeatedly proposed to instead model cognitive and emotional deficits that serve as core “ingredients” of mental dysfunction (*10*, *54*, *55*). We show the first working example of such a model, connecting a cognitive domain across species to a therapeutic intervention. Our example uses cognitive control, but other domains of function also change with DBS (*56–58*), and the same approach likely applies.

Our model has immediate clinical implications. For instance, impulsivity has been proposed as a mechanism of DBS’ therapeutic effect (e.g., by making a patient with OCD less sensitive to the “risk” of foregoing a ritual) (*34*, *59*). We show that impulsivity is dissociable from cognitive improvement, i.e. is likely a side effect. Similarly, our findings explain recent anatomic papers. Striatal DBS’ efficacy correlates with engagement of corticofugal fibers linking PFC to STN (*51*, *52*). That tract aligns with our mid-striatal target. In primates, this part of internal capsule carries corticofugal fibers from dorsal anterior cingulate (dACC), a structure heavily associated with cognitive control (*19*, *20*, *22*, *41*, *60*). We showed increased c-fos in PL, a potential homologue of dACC that projects through mid-striatum (*40*, *61*). Primate cognitive control and DBS’ effects also load onto lateral PFC (*19*, *20*, *28*), which has no known rodent homologue. Nevertheless, the alignment across other regions demonstrates the relevance of a cross-species model.

This and similar animal models should improve therapy development through new stimulation paradigms, by enabling more rapid screening. For instance, our RT-based measures of cognitive control can be tracked in real time to identify parameters that optimally improve them (*62*), which may enable rapid screening of novel paradigms such as temporally irregular (*14*) or plasticity-directed (*63*) stimulation. Similar screening in humans would require weeks of expensive hospitalization (*7*, *8*). Robust animal models could also improve DBS patient selection. Because mental illnesses are internally heterogeneous (*10*, *10*, *16*, *55*), neurostimulation of any given target will not work for most patients (*6*, *11*). Methods are emerging to identify patients whose symptoms arise from specific dysfunctions, including cognitive control (*64*). Pairing that identification with cognition-optimized neurostimulation could dramatically improve outcomes. Finally, our results enable causal neuroscience. Non-human primates have strong internal capsule and prefrontal homology to humans (*40*, *60*).

Striatal stimulation could manipulate specific sub-processes of cognitive control and uncover their neural substrates. Striatal stimulation altered drift rate across species, but boundary separation only in rats. This is due to the task structure and its effect on error rates (see Methods for elaboration). Rats require multiple trials to learn each Set-Shift rule (Figure 1). The human MSIT switched between high and low conflict (analogous to the Set Shift rules) every 1-2 trials. Rats can precommit to a strategy (following the light or going to a specific side) before they initiate a trial and can learn this strategy during a block. Figures 1-3 show that both processes occur. Strategic anticipation decreases the boundary separation, since less sensory evidence is needed to decide. Humans in MSIT can only process the sensory evidence once it is available, which loads onto the drift rate. Similarly, while rats frequently make errors, human performance on MSIT is almost always above 95% accuracy (*27*, *28*, *65*). Without a high error rate, it is impossible to detect boundary separation changes, because that DDM term captures speed-accuracy tradeoffs. In a different task design, e.g. with long runs of high or low conflict trials, humans would likely show learning and DBS would likely alter a boundary separation term.

Neuro-behavioral correlations highlighted PL and IL, but we did not see strong differences in c-fos evoked by stimulation at different striatal sites (Figure S8B), despite their different behavioral effects. Anatomy suggests that ventral stimulation should preferentially evoke c-fos expression in mOFC (*50*, *60*, *61*), but mOFC c-fos was numerically higher for mid-striatal and dorsolateral stimulation. This discordance is explained by our stimulation paradigm. First, our relatively high current (300 µA) likely spreads well beyond the electrode tip, creating overlap between different targets’ electric fields. Second, we stimulated for an hour before sacrificing for c-fos. Corticofugal fibers and striatal neurons form recurrent interconnected loops (*60*, *66*). Stimulation might have propagated through these connections to impact cortex more broadly. Third, 130 Hz stimulation has complex excitatory/inhibitory effects (*9*, *28*, *53*). C-fos only reflects excitation, and thus may not fully reflect cortical engagement.

This study had specific limitations in the model species and task. We studied only male rats, obscuring any potential sex differences. However, given that no sex differences were observed in the corresponding human studies (*27*, *28*), we did not anticipate significant sex-related differences. We also used outbred rats, not a disease model. This could limit the translatability of results to humans with psychiatric illnesses. However, as noted above, it is not clear that any putative “model of” psychiatric illness reflects human pathophysiology (*15*, *16*). Rat models of transdiagnostic impairments, such as compulsivity, exhibit motor alterations that could confound our results (*67*, *68*).

Regarding the task, we used set shifting to measure of the broader construct of cognitive control. This is the most commonly reported measure of cognitive control in rodents. Other assays of cognitive flexibility, such as reversal learning, have less need to suppress a prepotent response (*69–71*). That suppression of competing rules/interference is critical –in the original paper, there was no RT effect of DBS in tasks without interference (*28*). Other tasks, such as recently proposed rodent flanker paradigms (*72*), may be useful for measuring the generalizability of our effects.

In summary, we provide the first complete example of mapping a circuit-directed intervention and its cognitive mechanisms across species, demonstrating the value of a transdiagnostic approach to animal models of psychiatric illnesses and interventions. Our findings open the door to clinical interventions guided both by rigorous phenotyping and objective markers of target engagement, paired with animal studies to develop novel therapeutic strategies. They similarly can inform the basic science of cognitive control and executive function, by demonstrating that causal manipulations can affect specific subcomponents of those processes. Collectively, these impacts bring us closer to robust and personalized therapies for severe psychiatric illnesses.

## Methods

### Study Overview

We sought to develop an animal model of the core effect in our prior clinical work (*27*, *28*): that 130 Hz stimulation of striatum and/or corticofugal fibers passing through striatum leads to faster response times (RTs) in a task that requires cognitive control, and that this improvement occurs on all trial types. We further sought to replicate the finding of (*27*) that the effect varies with the precise striatal region being stimulated, with the largest effects from stimulation slightly dorsal to the ventral striatum. We thus tested a Set-Shift task that, similar to human tasks, requires inhibition of a pre-potent response (see below). Electrical stimulation parameters were chosen as homologues of those used in prior work (*27–29*), and statistical testing of the primary outcome similarly followed those papers. Partitioning of the striatum into separable stimulation zones (see below) was based on theoretical suggestions of a dorsal/ventral and lateral/medial functional segregation (*21*, *73*, *74*), anatomic studies showing a specific topography of corticofugal fibers passing through different striatal sub-regions (*60*, *61*), and prior work suggesting cognitive benefits of stimulation in the mid-striatum (*42*, *43*). We performed secondary (unplanned) analyses through drift-diffusion and related models because those models are particularly well suited to dissecting decision making under response conflict (*49*, *75*).

### Animals

81 male Long-Evans rats (250-300 g) were obtained from Charles River Laboratories (Wilmington, MA) and housed under a 10:14 dark/light cycle (lights on at 07:00). Rats were first acclimated for 5-7 days in the animal colony room and, subsequently, were handled for four consecutive days for 5 min/day to familiarize them with the experimenters. During the behavioral task and stimulation protocols described below, rats were food restricted. For this, animals were restricted to 10 g of standard laboratory rat chow per day until body weight was reduced to 85-90% of the original (after 5-7 days of food restriction, approximately). At this time food was increased to 10-15 g per day and their weights were maintained at this level without further reduction until completion of the behavior/stimulation experiments. Additional food supplementation was provided on days when an animal failed to earn sufficient behavioral task rewards to maintain its caloric needs. All experiments were approved by the University of Minnesota Institutional Animal Care and Use Committee (protocols 1806-35990A and 2104-39021A) and complied with National Institutes of Health guidelines.

### Set Shifting task

Sessions were performed in standard operant chambers (25 × 29 × 25 cm; Coulbourn Instruments, Holliston, MA), enclosed inside sound-attenuating boxes (Med Associates, Chicago, IL). Behavioral chambers were equipped with three nose-poke holes on one wall. Nose pokes could be illuminated and had infrared sensors to detect head entries. A food dispenser delivered reward pellets (45 mg grain-based pellets, Bio-Serv, Flemington, NJ) to a feeder on the opposite wall. Software and an appropriate interface (GraphicState 4.0, Coulbourn Instruments) controlled the presentation and sequencing of stimuli. Behavior was recorded by video cameras mounted on the top of each unit.

The operant Set-Shift task (Figure 1A) was modified from (*44*, *76*) and described in detail in a previous study from our group (*77*). In brief, animals learned and then shifted their responses between two distinct perceptual discrimination rules, or dimensions: a cue-driven “Light” rule and a spatial “Side” rule. Rats were required to poke the illuminated middle nose-poke hole to initiate a trial. In all trials, one of the two peripheral nose-poke holes was then illuminated. The Light discrimination rule required the rats to poke the illuminated nose-poke hole, regardless of its spatial location. The Side rule required that the animals poke at a designated spatial location across trials (left or right), regardless of which one was illuminated. The Light rule is substantially easier to execute because it requires a simple response to a visually salient stimulus, and thus creates a pre-potent response tendency that must be overcome on Side trials (see Figure 1 B-D). Rats were reinforced with a single reward pellet for each correct response. After reaching the performance criterion of 5 consecutive correct choices, the rule was switched to the other dimension, requiring rats to shift their behavior to continue receiving rewards. The sequential trials on a single rule, regardless of correct/incorrect choice, were grouped into a Block (Figure 1A). No explicit cue was provided, besides the absence of expected reward, to signal rule changes. The task required the rats to reach the performance criterion eight consecutive times, resulting in seven consecutive shifts per test session. To reach test performance, animals were submitted to habituation (3 sessions), shaping (3 sessions), and training phases (8-12 sessions). These were as described in prior work (*77*). They then underwent 16 testing sessions (see “DBS-like electrical stimulation” below). To prevent issues related to task structure (predictability of the next trial), 4 different test schedules, with different Side/Light rule orders, were administered using a pseudo-random test order (ABCDDCBAABCDDCBA, where each letter represents the schedule for a given testing day). This order was the same for each rat, except that any sessions that could not be run (e.g., if there were an equipment malfunction or abnormal testing conditions) were added to the end of the order. Additional non-contingent trials, in which rewards were given independently of the rats’ choice, were presented at the beginning and end of each session to prevent animals from discriminating between test schedules. Also, the within-test rule sequences were counterbalanced so that all possible discrimination rules were presented with equal frequency.

### 5 Choice Serial Reaction Time Task (5-CSRTT)

Previous studies with animal models (*78*) and humans (*37*, *79*, *80*) suggest that striatal DBS can increase impulsivity, and increased impulsivity could explain the observed RT reduction from DBS. To test this possibility, we evaluated stimulation’s effects in the 5-choice serial reaction time task (5-CSRTT, Figure 1H), a test of sustained attention that is also used to measure impulsivity (*46*).

The 5-CSRTT used operant chambers similar to those used for Set-Shift (Coulbourn Instruments, Holliston, MA), modified with a 5 nose-poke wall (Figure 1H). This module had 5 nose poke holes equipped with lights and infrared sensors to detect head entries. A food dispenser and a food trough, equipped with infrared sensors capable of detecting head entries used for trial initiation, were positioned on the opposite wall. Software and an appropriate interface (GraphicState 4.0, Coulbourn Instruments or in-house Pybehave/OSCAR (*81*)) controlled the presentation and sequencing of stimuli. Video recording was identical to Set-Shift.

The 5-CSRTT required rats to initiate a trial by nose-poking the illuminated food trough at the back of the chamber, then wait 5 seconds before a brief flash of light appeared in one of the 5 nose-pokes on the opposite wall of the chamber. The flash of light appeared for 0.5 seconds (stimulus duration) and rats had an additional 5 seconds (limited hold) to nose poke the aperture that lit up. Rats were reinforced with a single reward pellet for each correct response. A 5-second timeout before the next trial could be initiated occurred regardless of response type. If rats nose-poked too early, nose-poked the wrong aperture, or failed to respond within 5 seconds, the house light extinguished briefly to signify failure, rats were not rewarded, and there was a timeout as above. These behaviors were scored as premature responses, incorrect responses, and omissions respectively.

The 5-CSRTT protocol was adapted with minimal change from Bari et al. (*46*). The rats were submitted to 3 days of habituation before beginning training and testing sessions. On the first day of habituation, reward pellets were placed in each nose poke and the food trough. Each rat was then placed in the operant conditioning chamber for a period of 15 minutes to acclimate to the chamber. On Day 2, each rat was placed in the chamber and rewarded for nose-poking the food trough. On the last day of habituation, each rat performed a modified version of the task in which it did not have to wait for the nose-pokes to illuminate and there was no penalty for wrong choices. Training sessions for the 5-CSRTT consisted of 12 stages of increasing difficulty (Table S15). Later stages had shorter stimulus durations and limited hold periods with longer initiation to stimulus delay. The final stage required an initiation to stimulus delay of 5 seconds, a stimulus duration of 0.5 seconds, and a limited hold of 5 seconds after the light illuminated. Rats were required to achieve an accuracy of at least 80%, with less than 20% omissions and at least 50 correct responses, to begin the stimulation experiment. To ensure advancement through the stages, rats were trained twice per day, but only the first session of each day was considered for advancement. Each rat learned the experiment at different rates (Table S16) and thus qualified for surgeries at different times.

Animals completed the training stages, then underwent surgery as below. After recovering, each animal resumed training on the final training stage until its behavior returned to baseline before beginning the primary stimulation experiment. Once it successfully completed this training stage, it advanced to stimulation testing. During the primary stimulation experiment, each rat only performed 5-CSRTT once daily, at around the same time each day.

### Surgery for stimulating electrodes

Rats were initially anesthetized with 3-5% isoflurane in an induction chamber and were mounted in a stereotaxic frame, then maintained under anesthesia with 0.5-3% isoflurane for the duration of the surgery. We verified a surgical depth of anesthesia by toe pinch. Bipolar twisted pair platinum-iridium insulated electrodes (MS333/8-BIU/SPC, Plastics One, Roanoke, VA) coated with Vybrant® DiI cell-labeling solution (Invitrogen, Eugene, OR) were placed bilaterally in one of four regions of interest. Prior work in humans suggests that stimulation’s effects on cognitive flexibility are spatially specific, with the greatest effects just dorsal to the ventral striatum/capsule (*27*). The homologous fiber bundles in rats, however, are diffusely interpenetrated throughout the striatal grey matter. They do show a topographic distribution, such that different subregions of the rat striatum contain projections and corticofugal fibers from different PFC components (*40*, *60*, *61*). Supporting this, studies of extinction learning found that electrical stimulation enhanced extinction – but only when delivered to a region just dorsal to the canonical ventral striatum (*42*, *43*). Considering these prior results, we planned to test the effects of stimulation at four sub-targets within the rat striatum (Figure 1E): mid-striatum (1.4 mm AP, ±2.0 mm ML, and − 6.0 mm DV), ventral striatum (1.3 mm AP, ±2.0 mm ML, and −6.7 mm DV), dorsomedial striatum (1.4 mm AP, ±1.8 mm ML, and −4.5 mm DV) or dorsolateral striatum (1.4 mm AP, ±3.4 mm ML, and −4.5 mm DV). Each of these sites contains both cell bodies of the accumbens/caudate/putamen and corticofugal axons that project to both thalamus and brainstem (*41*, *61*). Rats were randomly assigned to each implantation group (Figure S1A). Several small burr holes were drilled around the perimeter of the exposed skull surface to accept anchor screws. Once the electrodes were placed, ground wires were wrapped to a nearby screw. The electrodes were then fixed to the skull and screws with dental cement. After surgery, rats were returned to their home cages and allowed to recover for 1 to 2 weeks before any behavioral tests.

For animals performing the 5-CSRTT, we only implanted electrodes in the mid-striatal target, based on the results of the primary Set-Shift experiment.

### DBS-like electrical stimulation

Electrical stimulation designed to model deep brain stimulation (DBS) was performed either using a PC running a custom-made LabVIEW program (National Instruments, Austin, TX) connected to a NI USB-6343 BNC analog/digital interface unit (National Instruments), connected in turn to an analog stimulus isolator (model 2200, A-M Systems, Sequim, WA), or using a PC running an Arduino script connected to a StimJim dual-channel electrical stimulator ((*82*), assembled by Labmaker, Berlin, Germany). We verified that both setups produced equivalent output current and waveforms. Stimulation parameters for both devices were set at 0.3 mA, 130 Hz, bipolar, biphasic, charge-balanced square wave pulses with a pulse width of 50 μs per phase, totaling 100 μs per pulse. 130 Hz was the frequency used in our prior human experiments and is a typical human DBS frequency (*27*, *28*, *59*). To ensure that any behavioral changes were not due to acute sensory percepts from stimulation, we delivered stimulation for 60 minutes prior to the Set-Shift and 5-CSRTT sessions and during the entire behavior testing session, without interruption. This pre-stimulation was conducted in a small holding arena adjacent to the operant chamber. For sham stimulation sessions, animals were connected to the stimulation cables for 60 min pre-test and during behavior testing, but no stimulation was given. Animals were submitted to the stimulation and sham stimulation sessions on interleaved days, for both behavior assays.

In a separate cohort of animals (n=6 implanted, n=5 included, see Figures S1 and S4), we tested the effect of changing the stimulation frequency. Frequencies between 10 and 50 Hz are usually ineffective in common DBS indications (*83*); we chose 20 Hz as a midpoint within this range that has also been associated with symptom worsening (*84*). These animals completed a protocol identical to the primary Set-Shift/130 Hz experiment. 3 weeks later, they were submitted to 8 additional sessions (4 sessions on, 4 sessions off), but with stimulation at 20 Hz (all other parameters unchanged from above). This thus includes an independent replication of our primary experiment’s core finding.

Conceivably, RT changes might reflect more general locomotor changes, e.g. an overall increased propensity to move. To rule this out, video recordings obtained during the pre-stimulation period were used to assess acute locomotor effects. Animal tracking in freely-moving rats was performed using DeepLabCut 2.3 (*85*), which focused on 10 body points of interest. The neural network was trained with a total of 1,079 frames, extracted from 13 randomly selected videos, for 600,000 to 1,030,000 iterations. Subsequently, the behavioral data, including metrics such as distance traveled, immobility, and the average velocity when moving, with poses estimated by DeepLabCut, were analyzed using DLCAnalyzer (*86*), a collection of custom R scripts.

In the 5-CSRTT cohort, some rats displayed seizure-like behaviors once mid-stimulation was initiated. Because this might reflect subtle electrode misplacement (not seen in the Set-Shift cohort), and because this directly affected testing, we excluded these rats from the analysis entirely and did not run them through the full experiment (Figure S5A).

### Histology and immunostaining

Before implantation, electrodes were coated with a fluorescent dye (DiI, Vybrant Multicolor Cell-Labeling Kit, Invitrogen, MA, US) to mark and help identify the tracks. There is no evidence that this dye can cause additional tissue damage or modify neural function (*87*, *88*). At the end of the experiments, animals were submitted to a last stimulation session (130 Hz only), delivered for 60 min to induce immediate-early gene (c-Fos) activation consistent with that induced by the main stimulation protocol. This allowed us to map cortical regions that were likely activated by the electrical stimulation and thus that might mediate observed behavioral effects. A separate cohort of animals (n=5) was used as a negative control. These subjects were not implanted with electrodes, did not receive any stimulation, and did not undergo any behavioral tests.

Ninety minutes after the end of the stimulation, rats were deeply anesthetized (pre-anesthesia with isoflurane followed by Beuthanasia-D Special, 150 mg/kg), and then transcardially perfused with cold phosphate buffered saline (PBS) followed by 4% paraformaldehyde (PFA) in 0.1 M PBS solution. Brains were removed and kept in PFA 4% at 4°C for 24 h and subsequently changed to 30% sucrose in 0.1 m PBS for 48-72h. Next, brains were snap frozen and sectioned on a cryostat.

For c-Fos immunostaining, sections were washed three times in PBST (PBS with 0.2% Triton-X) and incubated in PBST+NGS (5% normal goat serum) for 1 h. Rabbit anti-c-Fos (1:5000; part number 5348, Cell Signaling Technology, Danvers, MA) was incubated overnight at 4 °C. The next day, sections were washed 3 times in PBST, followed by 2-h room temperature incubation in Alexa 647 donkey anti-rabbit antibody (part number 711-606-152, Jackson ImmunoResearch, West Grove, PA). In the concluding steps of the staining protocol, to confirm probe placement, 4’,6-diamidino-2-phenylindole (DAPI, 1:10000) was applied to brain sections (20 min), followed by a PBS wash. Successful placement was defined when the tips of the electrodes were within the boundaries of the dorsomedial, dorsolateral, mid or ventral striatum as previously defined based on our pre-surgical targeting. Sections were mounted on slides, using VectaShield mounting medium, and coverslipped. Images were obtained using an inverted fluorescence microscope (Keyence, Osaka, Japan). FIJI (*89*) and the Trainable Weka Segmentation plugin (*90*) were used for quantification of cells expressing c-Fos protein in 720 × 540 µm areas of cingulate (Cg1), infralimbic (IL), prelimbic (PL), medial orbitofrontal (mOFC), lateral orbitofrontal (lOFC), and ventral orbitofrontal cortices (vOFC), as well as in striatal areas adjacent to the implanted electrodes (VS, MS, DMS, DLS). Cortical regions were identified by overlaying the imaged slice on a standard brain atlas (*91*). We chose these cortical regions because they are well established as sending fibers through our stimulation targets (*61*) and our past work suggests that DBS may act by retrograde modulation of these cortical origins (*27*, *28*).

### Statistical analysis - design and power considerations

Primary behavioral analyses were conducted using R version 4.3.2 (*92*) and RStudio version 2023.12.0.369 (*93*). Prior to model fitting we assessed the distribution of each response variable using the fitdistrplus package (*94*); the best-fitting distributions were used in the corresponding models. Other model-based analyses used specialized software described below in the relevant sections.

The primary Set-Shift experiment design and analysis were pre-planned, with the four stimulation sites selected on the basis of prior theory/anatomic subdivisions of the rodent striatum (see Study Overview above) and the analysis method (see below) selected to match the generalized linear models (GLMs) used in prior human studies (*27*, *28*). Animal counts and number of sessions were predetermined by a power analysis. Because we report 4 behavioral outcomes from the same task (see below), we targeted 80% power at a Bonferroni-corrected alpha=0.0125. Because a GLM is functionally equivalent to a repeated-measures ANOVA, we used that power estimator within G*Power 3.1. Assuming 8 measurements per animal and 4 groups, the canonical medium effect size (f=0.25) required 24 animals (7 per group). To provide a safety factor, we empirically increased this to 8 animals per group and 16 measurements per animal. We then allocated and implanted at least 10 animals per primary group (Figure S1A) to ensure that we had sufficient power after animal exclusions for technical failures.

As described further below, we ran a second cohort of the Set-Shift experiment, targeting only mid-striatal stimulation based on the primary findings. This aimed to validate the initial result and to test a further control, namely different stimulation frequencies. Because this study involved fewer comparisons, the effect direction was known, and we had observed in the primary experiments that effect sizes were larger than anticipated, we empirically decreased this to a planned sample of 6 rats. As described in the main text, this sample replicated fully the result of the main experiment.

The 5-CSRTT sample size was pre-planned to be identical in size and protocol (8 completed animals, 16 total testing sessions) to the primary Set-Shift sample. We similarly planned to do this experiment only if the Set-Shift experiment identified a behaviorally effective stimulation site. The analysis strategy was pre-planned to use GLMs with the same fixed/random effects as Set-Shift, but with distributions appropriate to the variables at hand (e.g., Poisson for response counts instead of gamma for response times). Due to animals that did not learn the task or had off-target electrodes, we eventually implanted a total of 27 animals rather than the 10 initially planned (Figure S5).

The c-fos regions for analysis and neuro-behavioral correlation strategy against Set-Shift and 5-CSRTT behavior changes were similarly pre-planned. No power analysis was conducted for this component, as our emphasis was on adequate power for the behavioral effect, and we did not plan to perform inferential statistics. With the large number of planned assessment regions, behavioral outcomes, and stimulation sites, a well powered study would require hundreds of animals.

The reinforcement learning and drift diffusion modeling, analysis of inter-trial behavior, and commonality analysis across experiments were not pre-planned. They were developed once the results of the primary Set-Shift experiments were known.

### Statistical analysis - Set Shifting behavior

For the analysis of reaction time in Set-Shift, we used generalized linear mixed models (GLMMs) to account for the repeated measures design and non-Gaussian distribution of the data. We analyzed reaction times (RT), accuracy, total time to complete each day’s testing session (affected by both RT and errors), and total error count within each session. To maintain consistency with our prior work (*27*, *28*, *95*) we used GLMMs with identity link function and gamma distribution for reaction time and for total time taken to complete each day’s session. For error counts, we used GLMMs with a log link function and Negative Binomial distribution. For accuracy (error rate/probability), we used GLMMs with a logit link and binomial distribution. We removed omission trials (where the rat did not respond during the trial window) from all analyses.

The general model for these analyses was:

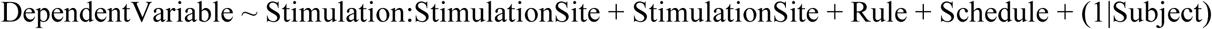

where Stimulation was coded 0/1 for stimulation on/off, StimulationSite was a categorical variable representing the implant site, Rule was Light or Side, and Schedule represented the specific block order schedule used on that testing day/session. Rule was included as a regressor for trial-level analyses (RT and accuracy), but not for session-level analyses (time to complete a testing session, total error count). Stimulation:StimulationSite mimics the fixed effect of Stimulation in our prior human papers, as does Rule. We added a test for a main effect of StimulationSite (i.e., the effect of the implant at that site when stimulation was off) because the electrodes used here are large relative to the anatomic site, and lesion effects might influence behavior. Schedule was a regressor of no interest and was treated as a fixed effect because random effects are generally not well estimated for factors with few levels. This model was chosen based on theoretical considerations and pre-specified hypotheses, not based on information criterion minimization or any other form of stepwise building. Model fitting was performed with the MASS (*96*) and lme4 (*97*) packages. Significance testing for all GLMM analyses was conducted by Wald Z-tests of the model parameters.

We applied a similar model structure and correction scheme for analysis of the second mid-striatal/Set-Shift cohort, which participated in the 20 Hz vs. 130 Hz experiment:

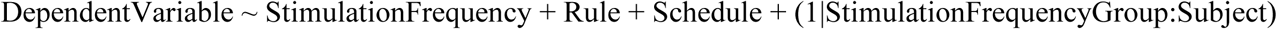

where StimulationFrequency is a categorical variable representing the frequency of stimulation, coded as 0 (off), 20, or 130 Hz. StimulationFrequencyGroup categorizes each subject’s sessions into subgroups based on the frequency of stimulation being delivered in a given experimental phase, either 20 Hz or 130 Hz. This is different from StimulationFrequency, because StimulationFrequencyGroup only takes the values 20/130, even on days when stimulation is off. We applied this additional random intercept because animals experienced the 20 Hz condition after the 130 Hz condition, i.e. after 16 more sessions of performing and learning the Set-Shift task. The additional intercept regresses out these practice effects, leaving only specific effects of 20 Hz stimulation.

We lumped the left and right Side rules together as a single rule on the basis of a similar GLM analysis demonstrating no statistical difference in Set Shift behavior between these two conditions; see Table S1.

### Statistical analysis - 5CSRTT behavior

For 5CSRTT, we used GLMMs to examine the number of premature responses, response accuracy, the number of omissions, and reaction time (RT). These are the standard read-outs of impulsivity in this task (*46*). To account for the nested nature of the data and individual differences among subjects, we included a random intercept for each subject in the model. For the analysis of premature responses, accuracy, and omissions, we employed GLMMs with a logit link function and a binomial distribution. For the analysis of RT, GLMMs with an identity link function and a gamma distribution were used, as they were with Set-Shift. Omission trials were excluded from the analyses of premature responses, accuracy, and RT to ensure that the data reflected completed trials. As with Set-Shift, we Bonferroni corrected the critical p-value to 0.0125 to account for the number of variables assessed (premature responses, RT, accuracy, and omissions).

The general structure of the statistical models was:

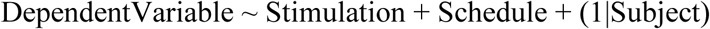

The dependent variable represented one of the behavioral measures, while “Stimulation” was coded as a binary variable (0 for stimulation off, 1 for stimulation on). The “Schedule” variable represents the specific block order schedule used on that testing day/session. There was no StimulationSite term because 5CSRTT animals were only implanted/stimulated at the mid-striatal target.

### Statistical analysis - Locomotor behavior and initiation delay

To evaluate potential locomotor and motivational changes as a cause of mid-striatal stimulation’s behavioral effects, we analyzed immobility, distance traveled, mean speed while animals were in the pre-stimulation arena, and delay time (time between when the cue light illuminated and when it was actually poked). We first assessed whether each variable had a relationship with RT by running generalized linear mixed effects models with a gamma distribution and identity link function.

The general structure of the statistical models was:

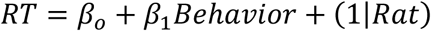

Secondly, we assessed whether each behavior was affected by stimulation by running generalized linear mixed effects models with a gamma distribution and identity link function for immobility and delay time and a linear mixed effects model for distance traveled and mean speed with a log transformation.

The general structure of the statistical models was:

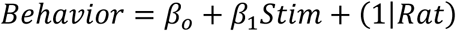

We performed this analysis only for animals implanted in the mid-striatum, after reviewing the results on the primary Set Shift analysis. We again applied a Bonferroni correction (critical p = 0.0125) to account for four dependent variables (immobility, distance traveled, speed, and delay time). There is no Schedule term because animals did not experience a specific behavioral schedule while in this pre-stimulation arena.

### Statistical analysis - Inter-trial behavior during Set-Shift

While observing video recordings as part of the above analyses, we noticed that rats would frequently nose poke ports during the inter-trial interval (ITI), despite such pokes being unrewarded. We hypothesized that these unrewarded pokes could be related to decision making processes and might reflect either positive or negative effects of stimulation. Specifically, unrewarded pokes might represent compulsive repetition of a prior response, regretful correction of that response post-error, or rehearsal for the next trial. If stimulation increases cognitive control, we might expect compulsive repetitions to decrease. Regretful corrections and pre-trial rehearsals, on the other hand, might increase, potentially reflecting proactive cognitive control allocation. During Side blocks in particular, the current trial’s outcome is 100% informative of the port that will be correct on the next trial, and thus rats could precommit to a response even before the cue. Extra nose pokes could add that precommitment, by setting up a response bias for the next trial. There is evidence that rats use such precommitment, in that they have slightly faster RTs on Side blocks (see Results, Figure 1C). Thus, stimulation that improves cognitive control should increase the probability of poking the port that will be correct on the following trial (and in many cases, that also would have been correct on the preceding trial).

We considered these hypotheses and potential influences of stimulation in a follow-up analysis for the mid-striatal cohort. Based on inspection of event densities (see Results, Figure 2), ITI behavior was divided based on poke location (left, middle, and right ports) and time (early: first two seconds, middle: middle three seconds, and late: last two seconds).

We first examined whether ITI behavior was related to performance and choices on the prior trial. For this analysis, we emphasized pokes in the early and late periods, with the early period capturing reactions to results of the previous choice (e.g., corrective or compulsive repetition) and the latter potentially reflecting practice for the upcoming trial. To evaluate if rats displayed corrective or rehearsal behaviors at the beginning or end of the ITI respectively, we used two Poisson GLMMs with an identity link:

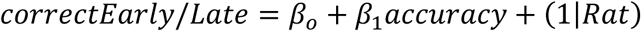

where correctEarly and correctLate are counts of the number of pokes in the previous trial’s correct port in the beginning or end of the ITI, while accuracy is an indicator if the rat responded correctly on the prior trial. Corrective behavior would yield a positive and negative for correctEarly (a tendency to nose-poke in the opposite port from the prior trial’s choice, but only when inaccurate), and rehearsal would yield a positive for correctLate (repeating the prior trial’s choice only when accurate on that trial).

Secondarily, because rats might have poked both ports but shown a bias towards one, we considered an additional model that treated ITI pokes as a continuous variable. This used binomial GLMMs with a logit link with separate models following correct or incorrect trials:

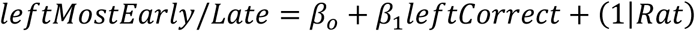

where leftMostEarly and leftMostLate are indicators for whether the left port was entered more often in the beginning or end of the ITI while leftCorrect is an indicator for whether the left port was the correct choice on the prior trial. The interpretation of follows the preceding model.

Third, we tested whether behavior during the ITI actually did influence behavior on the subsequent trial. This analysis used only late ITI pokes, which seemed most likely to influence behavior that occurred a few seconds later. To evaluate whether ITI behavior could predict choices in the subsequent trial we used a binomial GLMM with a logit link with separate models for trials where the light was/was not chosen:

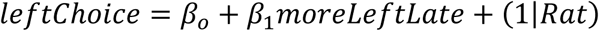

where leftChoice and moreLeftITI are indicators for whether the rat chose the left port during the subsequent trial and the left port more frequently during the last 2 seconds of the ITI respectively. Precommitment would be reflected in a positive and a negative (biasing the rat towards the right port by default and when there were fewer left port pokes in the late ITI). The separate models for when the light was/was not chosen capture the rat’s belief state regarding the current rule. If the rat currently believes itself to be in a Light rule block, then precommitment pokes should have little effect, because it will be attending to the cue light, and thus the effect of leftPokeLate should be attenuated in the light-chosen model.

Fourth, we tested whether ITI behavior could predict reaction times in the subsequent trial (and thus reflect a proactive cognitive control allocation reducing RT). We considered a possible confound, namely motivation. ITI pokes might represent precommitment to an upcoming response, but they might also reflect a general level of alertness or motivation to complete the trial and earn reward. These scenarios could be distinguished by which port is poked. Motivation should lead to increased pokes to the middle port, which is used to initiate a trial but is not a valid response. Precommitment, on the other hand, should lead to increased pokes in a port that the rat will also choose once the trial begins. To discriminate these two possibilities, and to test whether ITI pokes might be an underlying mediator of RT improvement, we used a gamma GLMM with an identity link:

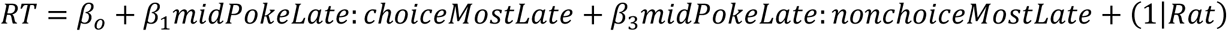

where RT is on the post-ITI trial and midPokeLate, choiceMostLate, and nonchoiceMostLate are indicators. midPokeLate is 1 if, in the last 2 seconds of the ITI, the rat made any middle port pokes. choiceMostLate and nonchoiceMostLate are mutually exclusive indicators capturing whether the rat’s pokes during the late ITI were more in the port matching the subsequent-trial choice (e.g., poke left during ITI and then choose left during the trial) or, conversely, the port not matching that subsequent choice.

Precommitment would be reflected in a positive or non-significant coefficient for nonchoiceMostLate (slower when pre-commiting to a port not subsequently chosen) and a negative coefficient for choiceMostLate (faster when pre-commiting to a port subsequently chosen). Motivation alone would be reflected, conversely, in a positive coefficient for midPokeLate, without substantial loading onto choiceMostLate or nonchoiceMostLate. The interaction between midPokeLate and choiceMostLate accounts for the possibility of rehearsal-like sequences of combined mid and side pokes having an additive effect on reaction times.

Lastly, the influence of electrical stimulation on late ITI pokes (choiceMostLate, nonchoiceMostLate, and midPokeLate) was evaluated via a set of binomial GLMMs with logit link functions:

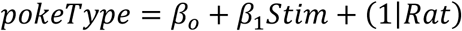

We ran these models only for animals implanted at the mid-striatal target. This enabled us to identify whether one or more specific ITI behaviors changed with mid-striatal stimulation, which in turn enabled the mechanistic analyses described in the following sections.

For these models, we did not correct critical values, as these were unplanned exploratory analyses to discover potential mechanisms.

### Statistical analysis - Reinforcement learning-drift diffusion model of Set-Shift behavior

To further dissect observed stimulation effects (see Results), we fit a series of computational models to the sequence of choices and reaction times made in each session. Traditionally, drift-diffusion models (DDMs) are the standard approach for such problems because they can explicitly predict reaction times and choices as a function of higher-level decision-making parameters (*49*). However, the choices rats favor in the Set-Shift task evolve across time as they receive rewards for choosing options consistent with the correct rule. Standard DDMs cannot account for these trial-to-trial changes in choice valuation. In contrast, Reinforcement Learning (RL) models are very effective for determining how reward history can be used to determine choice values (*48*). However, standard RL models do not predict reaction times. We addressed this dichotomy by instead using a family of combination reinforcement learning-drift diffusion models (RL-DDMs) which modify DDM parameters trial-to-trial based on RL valuation (*47*). On each trial, the RLDDM begins by assigning learned values to each port according to the location and illumination. In the example (Results, Figure 3), the left side is illuminated making its total value the sum of the values for the left side and light uniquely (*98*). The difference in values between the ports is then leveraged to make a choice based on a drift-diffusion process. The difference in total value informs the drift rate (rate of evidence accumulation towards a bound) while the difference in value between the sides alone informs the bias (initial evidence favoring one decision over the other). Trials where the value difference is high are likely to have shorter reaction times and more consistent choices while those with low value difference have longer reaction times and more variable choices. Once a port has been chosen, the values of the sides and light are updated based on a reinforcement learning process. The features of the chosen port are updated based on the prediction error with a non-linear scaling to diminish small errors and accentuate large ones. The value of the unchosen option decays according to a forgetfulness factor (*99*). These steps will repeat until the task is complete. The full set of equations for the model is as follows:

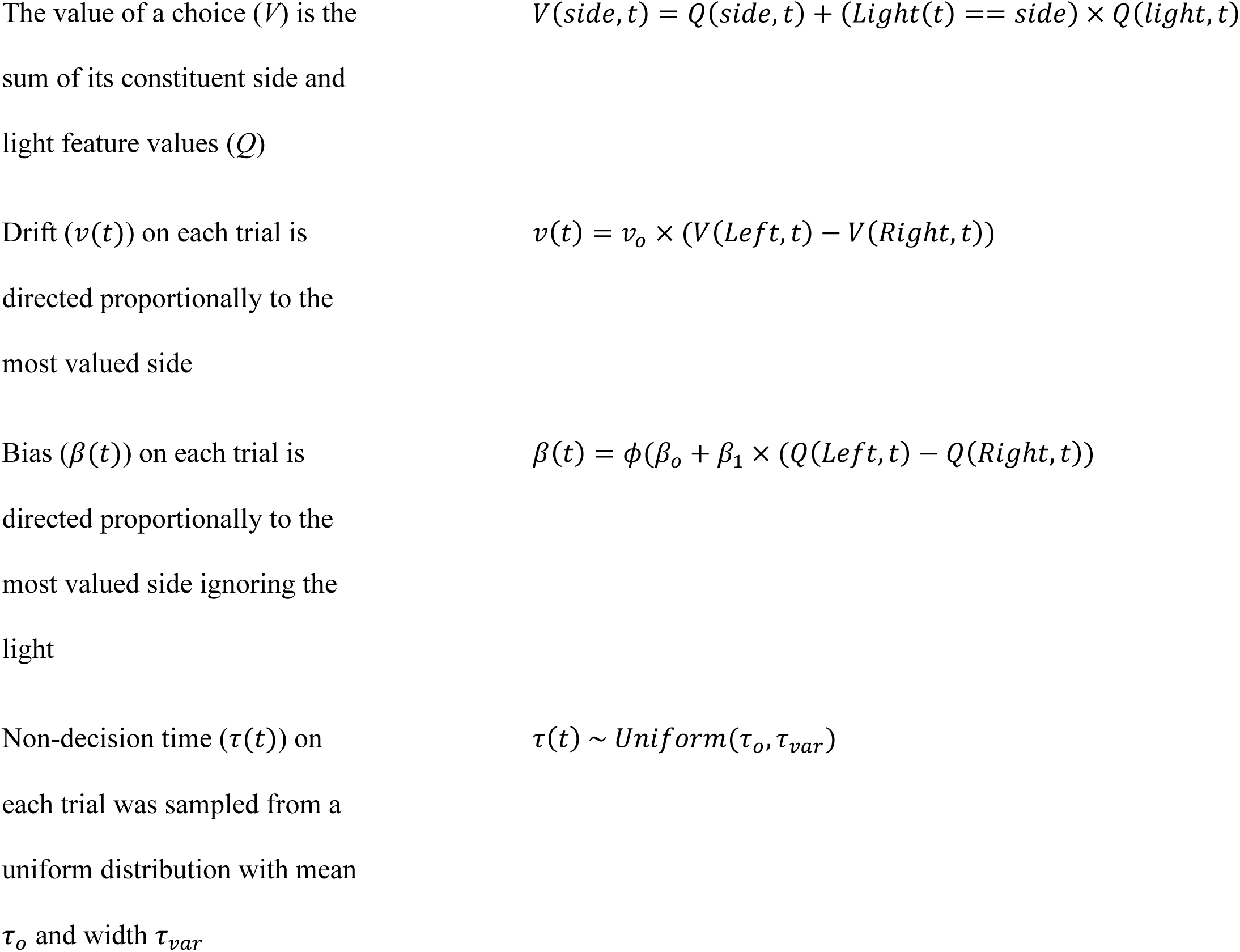
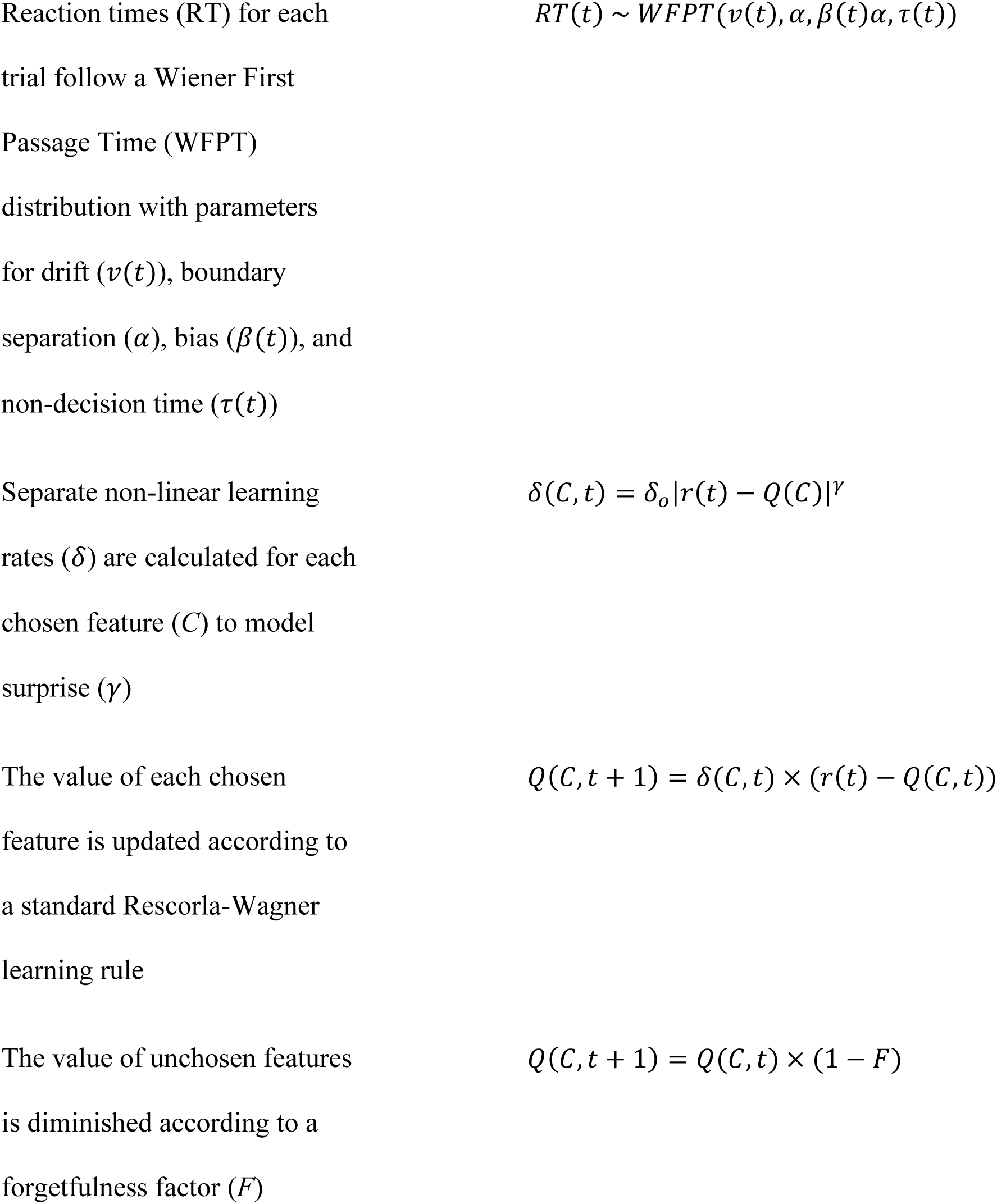

The baseline model has 8 parameters in total: initial values for the side and light features, drift rate scaling (*v_o_*), baseline bias (β_*o*_), non-decision time (*τ*_*o*_), non-decision time variability (*τ_var_*), boundary separation (*α*), and baseline learning rate (*δ*_*o*_). We considered three additional mechanisms namely, bias scaling (β_1_), surprise (*γ*), and forgetfulness (*F*), and fit different combinations of them for a total of 8 models. We chose the best model from these 8 by an information criterion process (see below). For all models, parameters were estimated using a Bayesian hierarchical modeling approach with levels for the group and individual subjects (see Figure S6 for schematic). Parameters for the initial values of the side and light features, baseline bias, and non-decision time variability were only estimated at the group level. All parameters estimated at the subject level had additional regressors for a fixed effect on stimulation-active sessions. All models were fit using the HDDM Python package (*100*). 2,000 samples with 1,000 burn-in samples were drawn from four MCMC chains (4000 total samples) and the model convergence was assessed by testing whether the Gelman-Rubin statistic (*101*) was <1.1. The predictive accuracy of the models were compared using Deviance Information Criterion (DIC). The best model according to DIC was further evaluated using posterior predictive checks to assess its ability to account for key features of the data. Simulated set-shift data was generated for each posterior sample using custom MATLAB code. For each of 4000 posterior draws, we used the estimated model parameters from that posterior draw (both group-level and subject-specific). We presented the RLDDM with the block sequence actually experienced by each rat, then allowed the DDM component of the model to generate a choice through a biased random walk (drift). Based on the outcome of that choice, we updated the RL parameters and stepped the model forward in time. We ran these simulations for both stimulated and non-stimulated sessions, applying the effects of stimulation as predicted by the model terms. Posterior predictive variables/checks included the reaction time distributions, error distributions, change in reaction times with stimulation, choice-value curves, and reaction time-value curves. These posterior checks depended on computing the choice that the model or the rat would make as a function of the value of the available options. Because those value estimates were not directly known for the rats, we estimated them through further simulations. We presented the RL component of the model with the choices actually made by the rat on each trial, and updated the terms within the model (Q and V) based on the received vs. model-predicted reward.

To determine the presence of a stimulation effect on model parameters, we assessed the group level distributions for the stimulation-specific parameters via two Bayesian statistics: the probability of direction (PD) and region of practical equivalence (ROPE) (*102*). PD refers to the proportion of the parameter distribution greater than 0 with a value above 0.5 (indicating a general increase in the parameter) and vice versa for values below 0.5. ROPE refers to the fraction of the density of the parameter distribution that lies within a region that is practically equivalent to a null-effect. To define our ROPE, we normalized our parameter estimates by dividing the group estimate by the pooled standard deviation parameters for the baseline parameter and stimulation-specific parameter. Using this normalized representation, we set our ROPE to be the region between +/- 0.1. PD is used as an indication for the existence/direction of an effect while ROPE establishes significance.

We additionally assessed the degree to which three model parameters (boundary separation, drift rate scaling, and bias scaling) could fully explain the observed decrease in reaction time with stimulation either uniquely or in common. Simulated set-shift data was generated by systematically removing the stimulation-specific effects for each parameter for a total of 8 different models each with 4000 posterior draws (simulated trials). The change in reaction time with stimulation active for each posterior draw was then compared across models relative to the empirical result using a commonality analysis (*103*, *104*). Note that because these models are not in any way meant to predict the behavior of individual rats, we did not perform the train/validate/test split that would be typical for a predictive regression model. The sole purpose of the RLDDM is to attempt to computationally explain the observed RT changes.

### Statistical analysis - Neurobehavioral correlations

Having identified manifest behavior changes and computational factors underlying those changes, we assessed which brain regions might be involved in both behavioral and computational effects of stimulation. As noted above, to identify physiological underpinnings of stimulation, we assessed c-fos counts in prefrontal regions known to be anatomically connected to the mid-striatal stimulation site and theorized to mediate stimulation’s cognitive effects (*11*, *51*, *52*, *59*), and in regions immediately neighboring the stimulation sites. We did not statistically compare counts between conditions/regions since the study was underpowered for this analysis.

We examined possible correlations between the effect of stimulation on behavior and c-fos expression. We limited this analysis to rats implanted in the mid-striatum (both cohorts), since that target showed the significant behavioral effect (see Results). Using a standard linear regression, we correlated the stimulation-induced change in per-subject boundary separation, drift rate, and bias with c-fos expression in each of the counted regions. Separate regressions were computed for each posterior draw and used to compute an average correlation and 95% highest density interval. We did the same for the change in manifest RT and estimated a confidence interval using a bootstrapped estimate with 4000 samples. Outlier c-fos counts were removed from these correlations based on the *rmoutliers* function in MATLAB with default arguments. We did not calculate statistical tests for this analysis, as the large number of correlations performed, without any pre-specified hypotheses, would likely yield false positives. We instead report the average correlation coefficient across the posterior draws descriptively, as a hypothesis-generating analysis.

### Statistical analysis - Drift diffusion modeling of prior papers’ human data

To further validate the conclusions of the drift-diffusion models described above, and to demonstrate the value of an animal model in providing mechanistic explanations, we attempted analogous modeling on the human participant data that inspired this work (*27*). Those data are publicly available, and all details on data collection, human participants protections, ethical approvals, and related topics are in the cited paper. Briefly, however, this dataset comprises trial-to-trial response times (RTs) and accuracy from participants performing the Multi Source Interference Task (MSIT; Figure 4A), a task intended to probe cognitive control. MSIT leverages classic sources of cognitive conflict/interference, such as the Flanker and Simon effects, both of which require cognitive control to resolve. Participants performing MSIT received stimulation to dorsal and/or ventral internal capsule while performing MSIT, at the same frequency (130 Hz) and roughly comparable amplitudes (usually 2 mA) to the parameters used here in rats. Importantly, the “dorsal” sites in that study were usually ventral to the head of the caudate nucleus (see Supplementary Figure 10 of (*27*)) implying that they corresponded more to the “Mid” than to the “Dorsomedial” site in our rats (see (*40*, *60*) for a more detailed anatomic discussion). Therefore, we focused our modeling on five participants who performed MSIT with and without “dorsal” capsule stimulation. We sought to test whether our primary rat finding (an increased drift rate) was also present in the original human data. Because of the task structure, two critical aspects of the modeling were different between rat and human. First, the rat Set-Shift task involves learning an explicit rule that changes across time, i.e. there is a component of reinforcement learning (leading to the RLDDM described above). In contrast, MSIT presents human participants with easy/hard (low/high conflict) trials in pseudorandom sequence, specifically to prevent response set learning. Thus, MSIT is best modeled using a basic DDM without the RL component. Second, rats make a large number of errors during Set-Shift (see Results, Figure 1) whereas humans make almost no errors on MSIT (*27*, *28*, *65*). Estimating changes in boundary separation requires a reasonable error rate that varies among testing conditions/trial types, because wider bounds tend to reduce errors and tighter bounds tend to increase them (*49*). Thus, although our DDM for MSIT includes a boundary term, we expected it not to vary with stimulation, essentially because of a floor effect.

We fit a basic DDM (Results, Figure 4B) to the RT and accuracy data (total of 1541 trials) using the same HDDM Python package used for the rat analyses (*100*). 3,000 samples with 2,000 burn-in samples were drawn from four MCMC chains and the model convergence was assessed by testing whether the Gelman-Rubin statistic (*101*) was <1.1. To test our hypothesis that stimulation would specifically increase the drift rate, we compared 4 models, including 1) DDM where drift rate (*V*) and boundary separation (*a*) parameters differ between stimulation and non-stimulation blocks, 2) DDM where stimulation only affects *V*, but not *a*, 3) DDM where stimulation only affects *a*, but not *V*, and 4) DDM model where both *V* and *a* do not differ between stimulation and non-stimulation blocks (no stimulation effect on parameters). We selected between these models based on minimization of the DIC, as we did for the rat modeling. The best model according to DIC was further evaluated using posterior predictive checks, again as was done for rat data. We performed posterior predictive checks on RTs using functions of the HDDM toolbox. A total of 770,500 simulated trials (1541 trials per simulation * 500 simulations) was sampled based on parameters drawn from the full posterior distribution of parameter estimates. We then compared the sampled RT distributions to the observed RT distribution in each trial type.

To further test the conclusions of this human DDM analysis (see Results), we fit the winning model (stimulation effect on drift rate *V*) to another dataset. Specifically, we used the patients from our original report of improved cognitive control from striatum/capsule stimulation (*28*). We fit the DDM to 14 participants’ RT and accuracy data (3944 trials; 3,000 samples with 2,000 burn-in samples from four MCMC chains). We performed the same comparisons as for the primary analysis on the data from (*27*).

## Funding

This work was supported by the OneMind Institute, the Minnesota Medical Discovery Team on Addictions, the MnDRIVE Brain Conditions initiative, the Tourette Association of America, and the US National Institutes of Health (R01NS120851, R01MH124687). ARO was supported by the São Paulo Research Foundation (FAPESP, Brazil—2017/22473-9. EDvR was supported by a National Science Foundation Graduate Research Fellowship under award number 2237827. The opinions herein are fully those of the authors and do not represent the positions of any funding body.

## Author Contributions

Conceptualization: ASW, AER

Data curation: AER, EDvR, MEM, EMS, AW

Formal analysis: AER, EDvR, JK, AA

Funding acquisition: ASW

Investigation: AER, MEM, EMS, AW, EH, KS, AA, DC, M-C L, ARO, GS, NS

Methodology: ASW, AER, EDvR, MEM, EMS Project administration, supervision: ASW

Writing - original drafts: AER, EDvR, JK, MEM

Writing - review and editing: AER, EDvR, MEM, ASW

## Competing Interests

ASW is a named inventor on multiple patent applications related to neurostimulation and cognitive control, none of which directly references the information herein and none of which is licensed to any entity.

## Data and Code Availability

Datafiles and analysis code will be made available via our laboratory GitHub repositories at the time of publication.

## List of Supplementary Materials

Figs. S1 to S10

Tables S1 to S26

## Notes

### Summary of Updates

Include missing sections/equations from the methods section due to file conversion.

