## supplemental_material for "Cross-species modeling and enhancement of cognitive control with striatal brain stimulation"

**The PDF file includes:**

Figs. S1 to S10
Tables S1 to S26

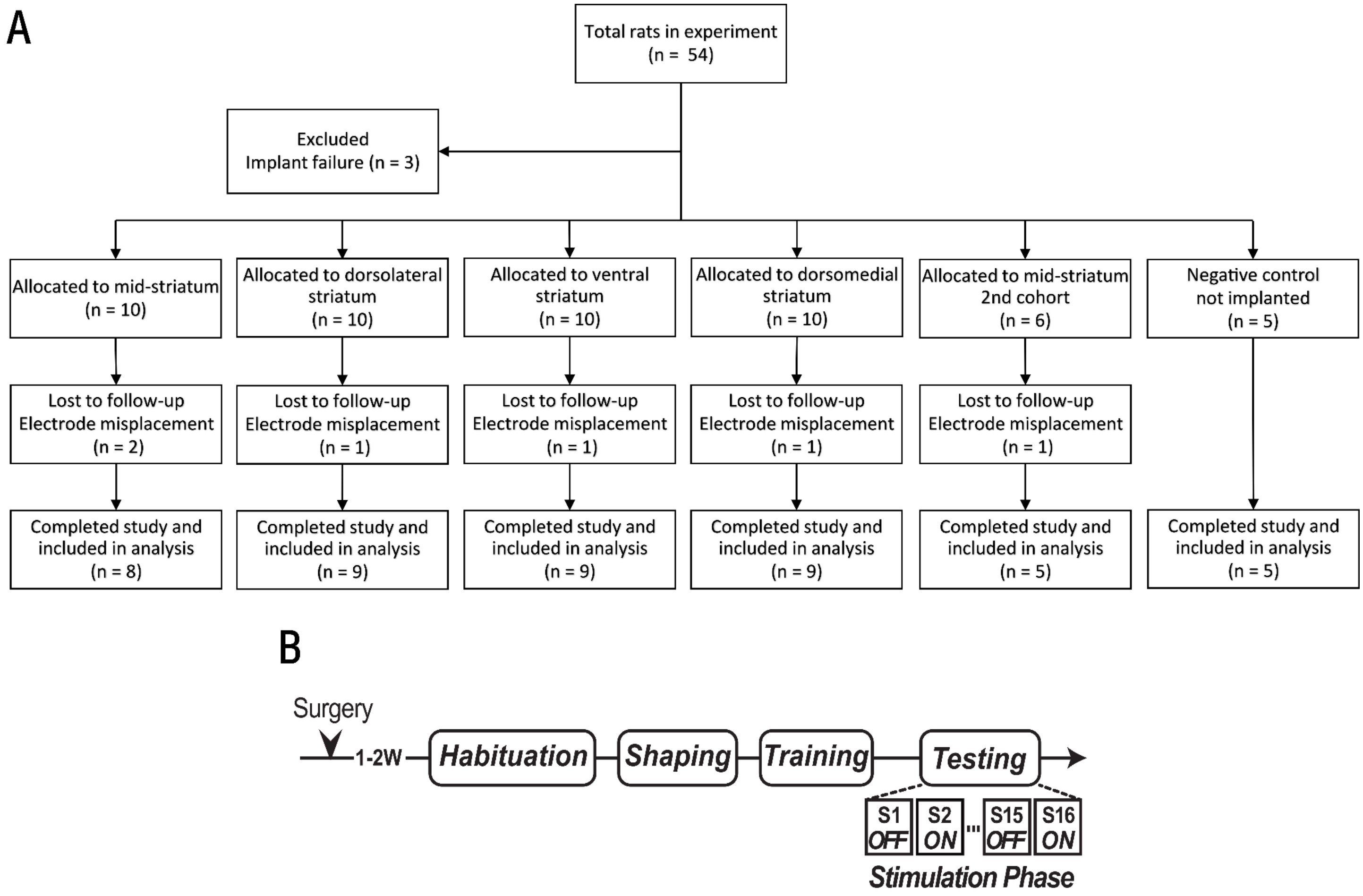


**Fig. S1:** Additional experimental details for primary Set-Shift experiment.
**(A)** CONSORT animal flow diagram for primary and secondary stimulation experiments. The second cohort of mid-striatal animals, as described in the Methods, was deliberately chosen as smaller based on the observed large effect size and because it was specifically planned as a confirmatory experiment.
**(B)** Experimental schedule diagram. After implant and training, rats underwent 16 sessions of testing, divided into alternating days of mid-striatal 130 Hz stimulation and no stimulation (still connected to stimulator/cables). This illustrates the primary cohort. The 2nd cohort had an additional testing phase of 20 Hz stimulation, conducted 3 weeks after the primary 130 Hz experiment.

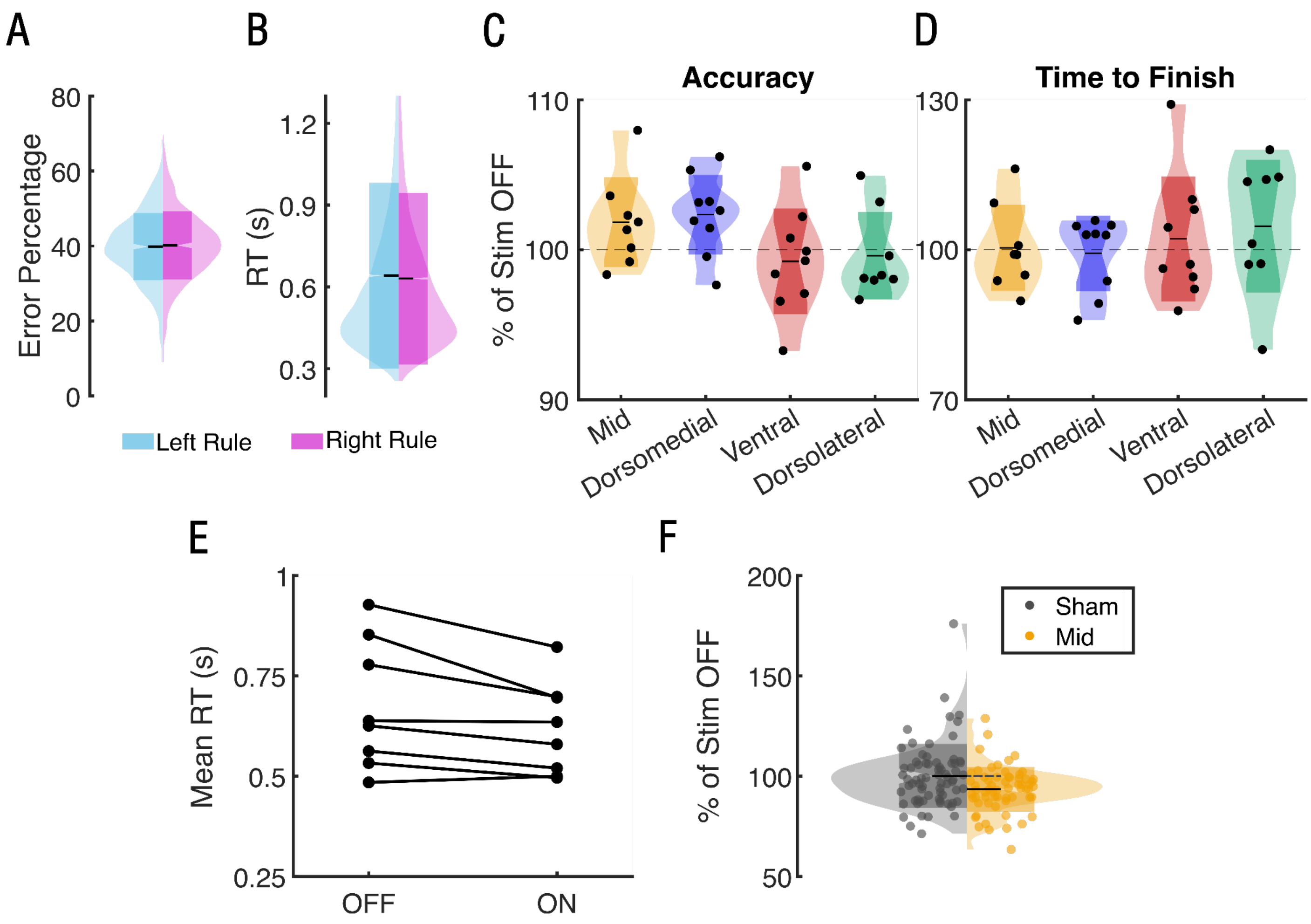


**Fig. S2:** Set-shift behavioral results and further outcomes of stimulation.
**(A)** Probability of error based on the current variant of the Side rule (left/right nose poke port). There was no significant difference between the two rules ($\beta$ = -0.006, p = 0.15), demonstrating that as a group, rats did not have a bias for a particular side of the chamber.
**(B)** RT based on the current variant of the Side rule (left/right), again showing no significant difference ($\beta$ = -0.033, p = 0.18).
**(C)** Distributions of accuracy as a function of stimulation type, expressed as a percentage of stimulation-OFF. Distributions are computed over rats; each dot shows the median for one rat. As with the total number of errors reported in the main text, there was no significant difference for any group (lowest p-value, p = 0.15 for Stim_ON:Site_dorsomedial), although mid and dorsomedial striatal stimulation did cause a non-significant improvement.
**(D)** Distributions of the total time to finish the task as a function of stimulation type, again showing no significant change for any stimulation location (p = 1.00 for all sites)
**(E)** Individual absolute change in RT for each rat receiving mid-striatal stimulation, expressed as the median RT over all stimulation ON or stimulation OFF testing sessions. For 6/8 rats, this shows a decrease.
**(F)** Distributions of RT for sham (gray) and mid-striatal (yellow) stimulation over sessions, relative to the median stimulation OFF value for each animal. Each dot shows an individual session. The dashed gray line extending into the stimulation side shows the line of no effect. 80.7% of the individual mid-striatal stimulation sessions fall below this line.

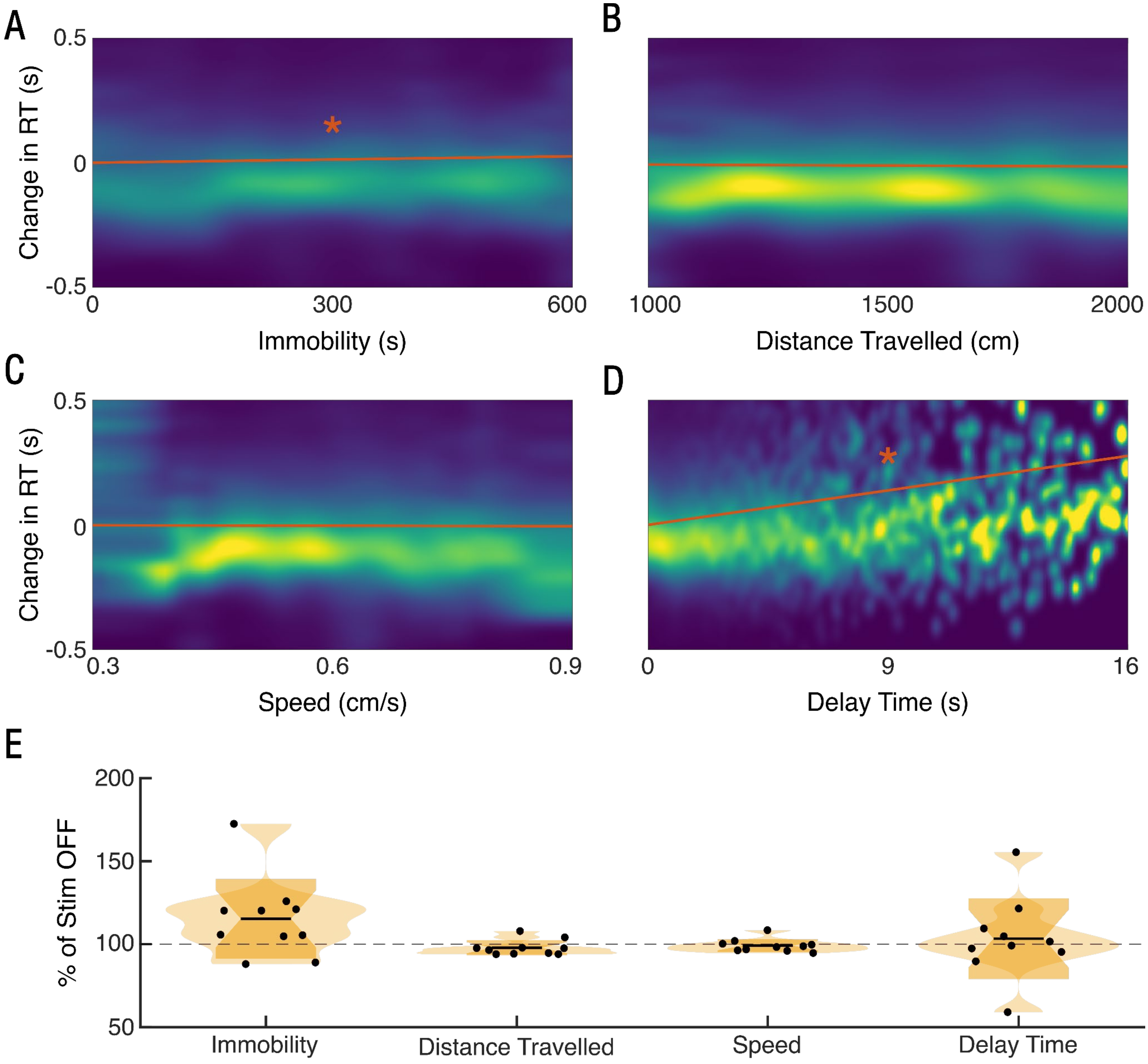


**Fig. S3:** Mid-striatal stimulation does not affect motor performance or motivation.

**(A-D)** 2D heatmaps of the kernel densities for RT as a function of motor behavior. At each point along the X-axis, the heatmap shows the distribution of RTs given that value of the corresponding motor variable. RTs are corrected by removing each rat's specific random intercept from the corresponding GLM (see "Locomotor behavior" in Methods). At each X-axis point, the distribution is normalized to have unit density for each column of the distribution to aid in comparison across motor performance and delay time values. The slope term from the corresponding GLM is indicated by the orange line overlaying each heatmap. Distance traveled, immobility, and speed were assessed for 50 min prior to each Set-Shift task run, in an open field. Delay time (panel D) was assessed during each Set-Shift task session. It is defined as the time between when the middle (trial initiation) port became illuminated and when the rat actually poked it. This

latency to initiate a trial is a measure of a rat's general motivation/drive to engage with the task. The degree of immobility (A) and delay time (D) were both significantly related to reaction times (see Supplementary Tables 6 & 9) while distance traveled (B) and speed (C) did not impact reaction times during the task.
**(E)** Change in motor variables with active mid-striatal stimulation, as a percentage of the corresponding behavior without stimulation. Values for individual animals are indicated by black dots. None of the behaviors were significantly affected by stimulation (all $p > 0.05$).

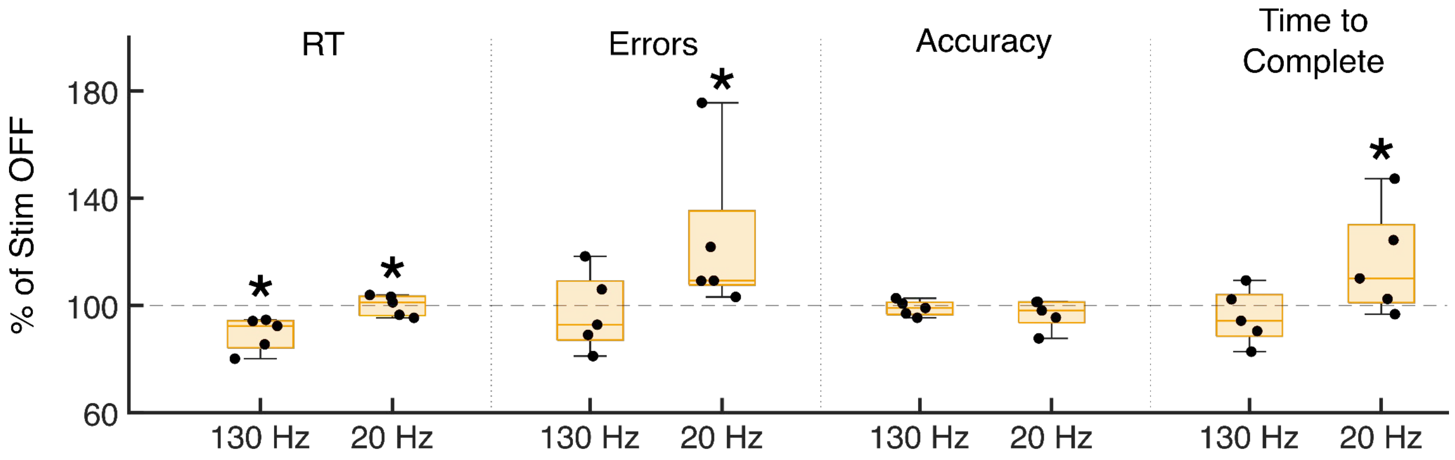


**Fig. S4:** Mid-striatal stimulation's effects in Set-Shift are frequency specific.

In a separate cohort of rats (n=5), 130 Hz interleaved ON-OFF sessions were performed similar to the original experiment. Three weeks later, the same rats underwent 20 Hz ON-OFF interleaved testing sessions (14871 trials across all testing sessions). 20 Hz stimulation worsened performance, with significant increases in RT ($\beta$ = 25 ms, p = 2.96e-4), errors (β=0.23, p=5.50e-4) and time to complete the overall session (estimate = 188 seconds, p < 0.017).

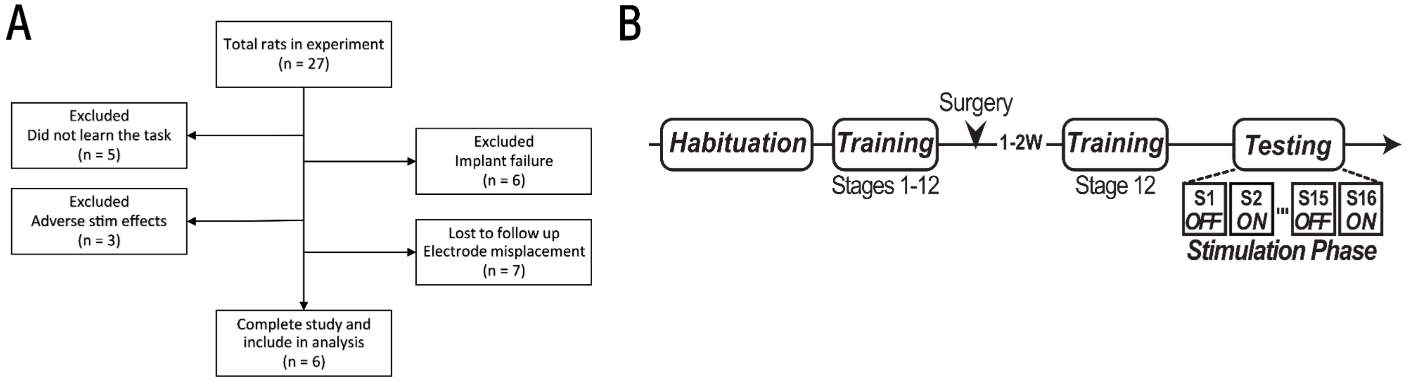


**Fig. S5:** Additional experimental details for 5-choice serial reaction time task (5CSRTT) experiment.

**(A)** CONSORT animal flow diagram. Implant failures were primarily head cap failures. Adverse stimulation effects were primarily seizure-like activity, marked by behavioral arrest and motor twitches.

**(B)** Experimental schedule diagram. After training and implant, plus training to return to criterion (see Supplementary Table 15), rats underwent 16 sessions of testing, divided into alternating days of mid-striatal 130 Hz stimulation and no stimulation (still connected to stimulator/cables).

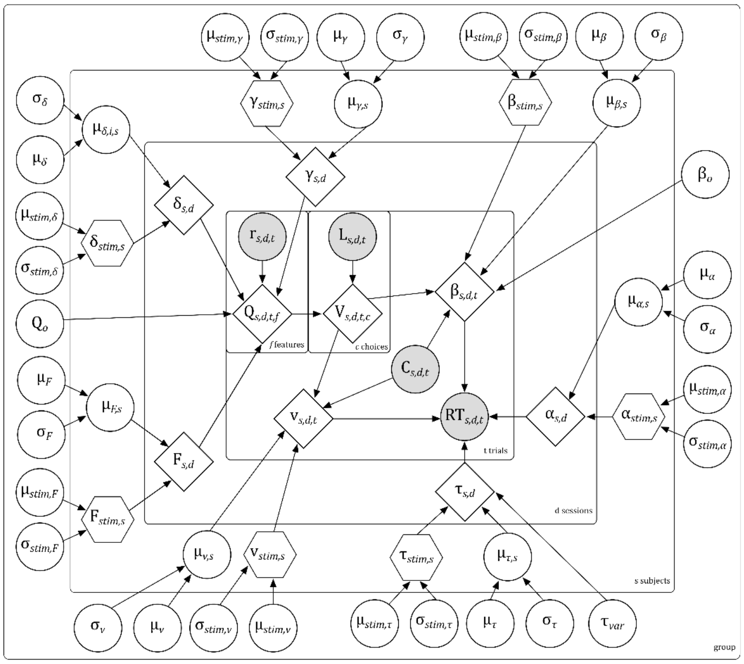


$\mu_\alpha$ ~ G(1.5,0.75)
$\mu_v$ ~ N(2,3)
$\mu_\tau$ ~ G(0.4,0.2)
$\mu_\beta$ ~ N(0,3)
$\mu_\delta$ ~ N(0.2,3)
$\mu_\gamma$ ~ N(0,3)
$\mu_F$ ~ N(-2.5,3)
$\mu_{stim,X}$ ~ N(0,3)
$\beta_o$ ~ N(0,3)
$Q_o$ ~ B(1,1)
$\tau_{var}$ ~ HN(0.3)

$\sigma_\alpha$ ~ HN(0.1)
$\sigma_v$ ~ HN(2)
$\sigma_\tau$ ~ HN(1)
$\sigma_\beta$ ~ HN(2)
$\sigma_\delta$ ~ U($10^{-10}$,10)
$\sigma_\gamma$ ~ U($10^{-10}$,10)
$\sigma_F$ ~ U($10^{-10}$,10)
$\sigma_{stim,X}$ ~ HN(2)

Model parameter
Stim parameter
Calculated parameter
Observed data

**Fig. S6:** Schematic of hierarchical reinforcement learning drift diffusion model, illustrating the model parameters (refer to main text Figure 3) and their nesting at the level of subjects, sessions, and trials.
Each node represents a parameter or variable used in the model. Shaded nodes represent observed variables, while non-shaded nodes represent latent variables. Squared nodes are deterministic parameters that are calculated as a function of other nodes. Non-shaded circular and hexagonal nodes are both fit parameters, with hexagonal nodes only influencing the model behavior when stimulation is active.

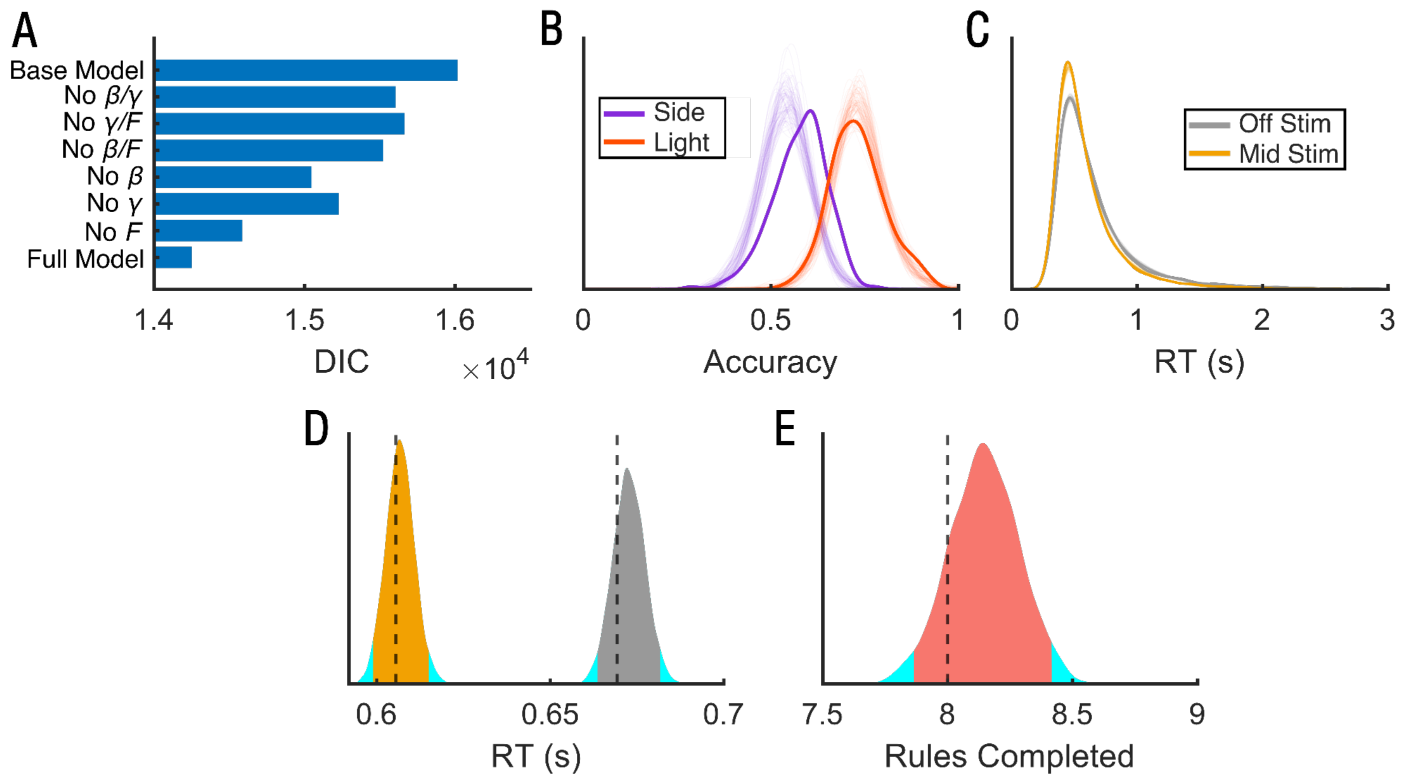


**Fig. S7:** Validation of Reinforcement Learning-Drift Diffusion Model
**(A)** Comparison of diffusion information criterion (DIC) between candidate models, with each bar showing the average DIC value over the four chains (1,000 samples each after burn in). The base model included animal-specific terms for drift rate, boundary separation, learning rate, and non-decision time along with group terms for overall bias, non-decision time variability, and initial RL values. Additional models included combinations of three more terms: forgetfulness (F), bias ($\beta$), and surprise ($\gamma$). The full model included all of these features and had the lowest DIC value (14251).
**(B)** Comparison of accuracy between side (purple) and light (orange) rule for simulated RLDDM and empirical rat data. Each sample from each chain was used to generate a distribution of accuracy values across sessions for each rule (transparent curves). The distribution for the empirical behavioral data is shown by the solid curve. The light rule accuracy was fully consistent with model predictions while the side rule accuracy was somewhat underestimated, although the distribution medians remained within 6.6%.
**(C)** Comparison of RTs between stimulation OFF (gray) and active mid-striatal stimulation (yellow) for simulated RLDDM and empirical rat data. Each sample from each chain was used to generate a distribution of reaction times for both conditions. The distribution for the empirical data is shown by the solid curve. Both ON and OFF distributions were fully consistent with model predictions, to the point that the simulation data curves completely overlap the empirical curves.
**(D)** Comparison of median RT between stimulation OFF (gray) and active mid-striatal stimulation (yellow) for simulated RLDDM and empirical rat data. Distributions are computed over all posterior draws, with the 95% highest density interval shown by the solid yellow or gray region. The median value for the rat data is shown by the dashed line. Median RT for both active stimulation and stimulation OFF was consistent between model simulations and rats (the empirical median was well within the simulated highest density interval).
**(E)** The total number of rules completed was consistent between rats and model simulations. Plotting conventions follow **(D)**.

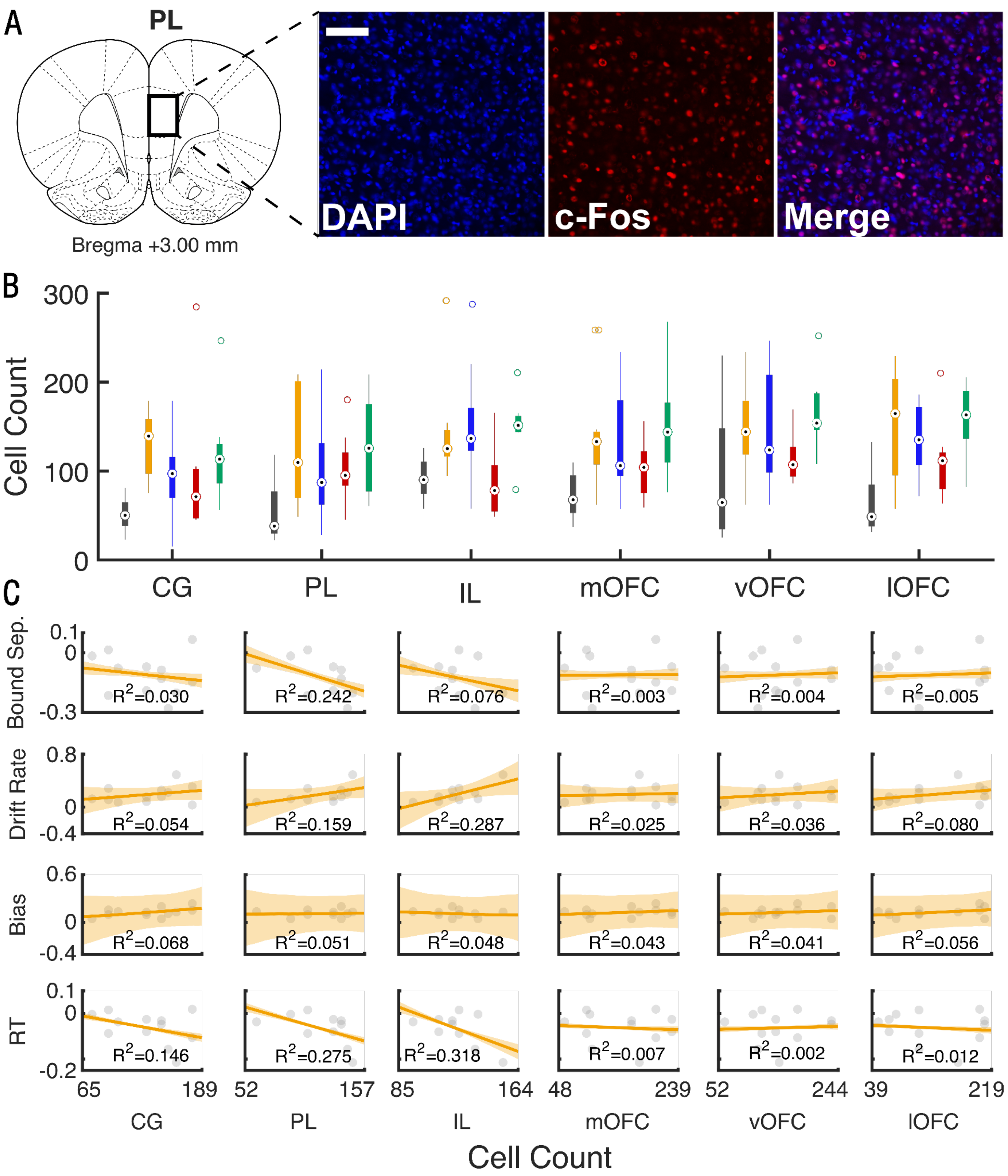


**Fig. S8:** Mid-striatal stimulation may exert its cognitive control effects through prefrontal modulation, with specific

cortical regions contributing to specific cognitive operations.
**(A)** Representative image of a field of view analyzed for c-fos staining in the PL after 1 hour of stimulation delivered to the mid-striatum. DAPI labeling (blue) and c-Fos immunostaining (red). Scale bar 100 μm.
**(B)** C-fos counts in ≥ 8 rats per stimulation site, two fields per rat. Circles show median and outliers, boxes and whiskers show 25-75th quantiles. Gray bars show rats that did not receive any stimulation before sacrifice (n = 5), colored bars show rats receiving stimulation in different striatal zones according to the color scheme in main text Figure 1. All stimulated rats received 1 hour of bilateral 130 Hz stimulation before sacrifice. Stimulation in almost every striatal sub-zone increased c-fos expression in all regions of prefrontal cortex, with limited preferential influence of any striatal zone on specific cortical regions.
**(C)** Correlation between changes in RL-DDM parameters reported in main text Figure 3 and c-fos expression in connected cortical regions, for mid-striatal stimulation only. The final row shows raw RT for comparison. Points show individual animals' RT or point estimate of model parameters. Lines show best fit, while shaded regions show the confidence interval of that fit based on the 4,000 posterior draws. We emphasize the three parameters that changed with stimulation and explained the primary RT change. Boundary separation decrease correlated modestly with PL modulation, and to a much lesser degree with modulation of IL. Drift rate increase, in contrast, correlated most strongly with IL modulation and less with PL. The manifest RT change correlated with PL, IL, and Cg c-fos, consistent with the main text finding that the RT improvement involves simultaneous changes in drift rate and boundary separation. We did not perform inferential statistics for these exploratory, non-planned analyses. Cg, cingulate; PL, prelimbic cortex; IL, infralimbic cortex; OFC, orbitofrontal cortex, with m=medial, v=ventral, l=lateral. Anatomic regions are as defined on plate 11 (Bregma +3.00 mm) of the 6th edition of the Paxinos and Watson atlas.

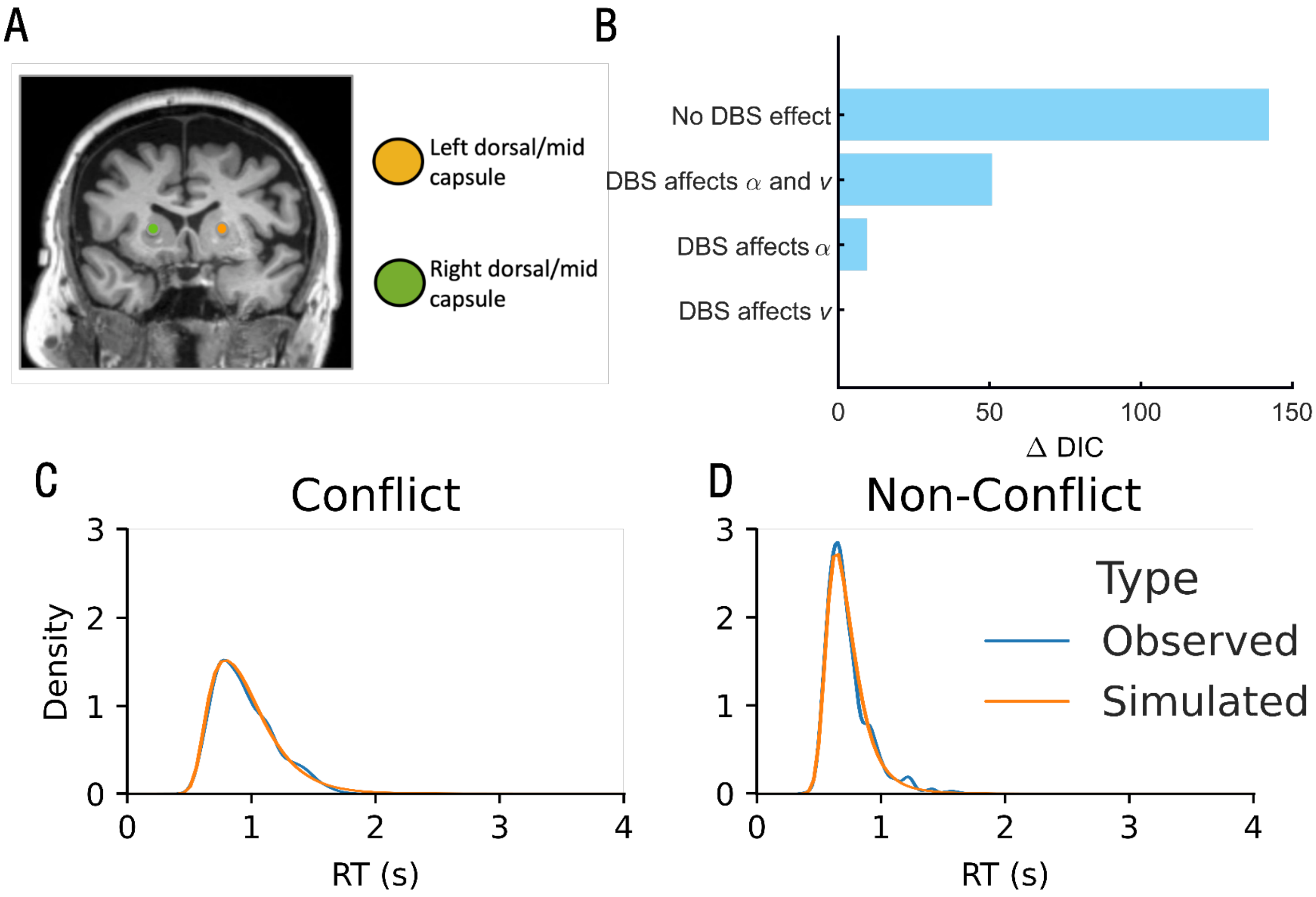


**Fig. S9:** Supplemental methods and modeling checks for drift-diffusion modeling on the prior human results of Basu et al. 2023.

**(A)** Schematic showing the locations of brain stimulation delivery, overlaid on an example patient's MRI. The original experiment involved stimulation at both ventral and dorsal/mid-internal capsule sites; we focused this analysis solely on patients stimulated more dorsally because this was behaviorally effective and mimicked the pattern seen in rats. Note that this "dorsal" stimulation is close to the "mid" site in rats; the equivalent of our "dorsomedial" rat site would be at or above the caudate.

**(B)** Model selection. We compared models where DBS-like stimulation had no effect on the model, vs. affecting drift rate ν, boundary separation α, or both. The model with stimulation affecting drift rate had the lowest DIC, as predicted based on rodent results and differences between the human MSIT and rat Set-Shift paradigms.

**(C)** Posterior predictive checks. Distributions show simulated and empirical RTs for 779 actual and 389,500 (779 trials per simulation * 500 simulations) simulated trials (Conflict) and 762 actual and 381,000 simulated trials (Non conflict). The empirical and simulated distributions almost exactly overlap.

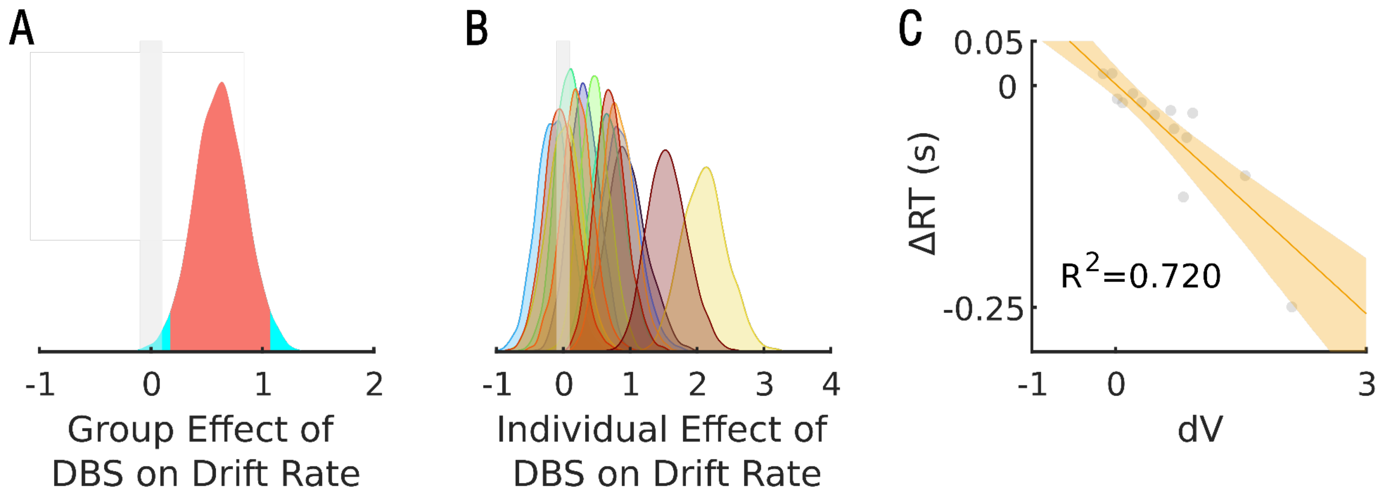


**Fig. S10:** Drift-diffusion modeling results from the prior human data of Widge et al. 2019, replicating the analysis of Figure S9 in an independent dataset.

**(A)** Distributions of stimulation effect on drift rate over 4000 posterior draws. The median of the distribution is indicated by the solid line, while the 95% highest density interval is in orange. The gray shaded area around zero represents the Region of Practical Equivalence (ROPE) for a null effect (effect size less than 0.1). Electrical stimulation substantially increased the drift rate (d=0.62, pd=0.996, 1.03% in ROPE).

**(B)** Individual distributions of drift rate changes, over 4000 posterior draws, on the same scale as **(C)**. In 7 of the 14 participants, stimulation substantially increased drift rate. Note that in contrast to Figure S9, in these participants, stimulation was not targeted to any specific striatal zone, but was left at clinical settings, which varied dramatically. As such, we would expect less consistent results than Figure S9 or the rodent model.

**(C)** Correlation between drift rate change and stimulation-induced RT change. Each dot represents one participant, specifically the RT change plotted against the point estimate of the drift rate change (median of the distributions shown in panel B). The solid line shows a line of best fit. The shaded region represents the 95% highest density interval of this line. We calculated this by running a separate correlation for each posterior draw (distributions in panel B) against the change in median RT.

**Table S1:** Regression coefficients for the task accuracy (proportion of correct responses) and reaction time (RT) in seconds for the two variants of the Side rule (left/right nose poke port) in the set-shifting experiments, for animals implanted in all sites. The coefficient for task accuracy is derived from a generalized linear model with a binomial distribution and logit link function. The coefficient for RT is derived from a generalized linear model with a gamma distribution and identity link function. Bonferroni correction applied for both coefficients.

There were no differences in accuracy or RT when the correct Side was right vs. left. We therefore collected these together as a single Side rule for all subsequent analyses. These data are also presented in Figure S2.

General formula: Variable ~ Correct_Side + (1|Subject)

Fixed effects coefficients (95% CIs):

| Outcome | Name | Coefficient | SE | tstat | DF | p.value | Adjusted.p |
|---|---|---|---|---|---|---|---|
| Error | (Intercept) | 0.386 | 0.013 | 28.774 | 43332 | 2.2e-16 | |
| | CorrectSide_Right | -0.033 | 0.020 | -1.699 | 43332 | 0.089 | 0.179 |
| Reaction time | (Intercept) | 0.641 | 0.002 | 292.35 | 43332 | 2.2e-16 | |
| | CorrectSide_Right | -0.006 | 0.003 | -1.790 | 43332 | 0.074 | 0.147 |

**Table S2:** Regression coefficients for reaction time (RT) in seconds in the set-shifting experiments, from a generalized linear mixed-effects model, with a gamma distribution and identity link function. Bonferroni adjustments are shown applied to the stimulation coefficients, which were our primary hypothesis and analytic focus. Mid-striatal stimulation improved RT, while dorsomedial stimulation marginally worsened it.

Formula: RT~ Stimulation:StimulationSite + StimulationSite + Rule + Schedule + (1|Subject)

Fixed effects coefficients (95% CIs)

| Name | Coefficient | SE | tstat | DF | p-value | Adjusted.p |
|---|---|---|---|---|---|---|
| (Intercept) | 0.575 | 0.052 | 11.084 | 71991 | 1.577e-28 | |
| Site_dorsolateral | 0.067 | 0.073 | 0.911 | 31 | 0.369 | |
| Site_dorsomedial | 0.135 | 0.073 | 1.837 | 31 | 0.076 | |
| Site_mid | 0.103 | 0.076 | 1.369 | 31 | 0.181 | |
| Rule_Side | -0.012 | 0.002 | -6.197 | 71991 | 5.778e-10 | |
| Schedule_2 | 0.004 | 0.003 | 1.564 | 71991 | 0.118 | |
| Schedule_3 | -0.004 | 0.003 | -1.401 | 71991 | 0.161 | |
| Schedule_4 | -0.003 | 0.003 | -0.941 | 71991 | 0.347 | |
| Stim_ON:Site_ventral | -0.002 | 0.004 | -0.476 | 71991 | 0.634 | 1.000 |
| Stim_ON:Site_dorsolateral | 0.008 | 0.004 | 1.839 | 71991 | 0.066 | 0.263 |
| Stim_ON:Site_dorsomedial | 0.018 | 0.004 | 4.382 | 71991 | 1.176e-05 | 4.704e-05 |
| Stim_ON:Site_mid | -0.034 | 0.004 | -8.624 | 71991 | 6.588e-18 | 2.635e-17 |

**Table S3:** Regression coefficients for task accuracy (proportion of correct responses) in the set-shifting experiments, from a generalized linear mixed-effects model, with a binomial distribution and logit link function. Bonferroni adjustments are shown applied only to the stimulation coefficients, which were our primary hypothesis and analytic focus.

Formula: Accuracy ~ Stimulation:StimulationSite + StimulationSite + Rule + Schedule + (1|Subject)

Fixed effects coefficients (95% CIs)

| **Name** | **Coefficient** | **SE** | **tstat** | **DF** | **p-value** | **Adjusted.p** |
|---|---|---|---|---|---|---|
| (Intercept) | 1.006 | 0.037 | 27.502 | 71991 | 1.000e-26 | |
| Site_dorsolateral | 0.117 | 0.045 | 2.601 | 31 | 0.014 | |
| Site_dorsomedial | 0.007 | 0.045 | 0.157 | 31 | 0.877 | |
| Site_mid | -0.103 | 0.045 | -2.285 | 31 | 0.029 | |
| Rule_Side | -0.637 | 0.017 | -38.385 | 71991 | 1.000e-26 | |
| Schedule_2 | -0.002 | 0.023 | -0.084 | 71991 | 0.933 | |
| Schedule_3 | -0.028 | 0.023 | -1.222 | 71991 | 0.222 | |
| Schedule_4 | 0.012 | 0.022 | 0.533 | 71991 | 0.594 | |
| Stim_ON:Site_ventral | -0.001 | 0.031 | -0.037 | 71991 | 0.970 | 1.000 |
| Stim_ON:Site_dorsolateral | -0.025 | 0.034 | -0.730 | 71991 | 0.465 | 1.000 |
| Stim_ON:Site_dorsomedial | 0.066 | 0.032 | 2.086 | 71991 | 0.037 | 0.148 |
| Stim_ON:Site_mid | 0.016 | 0.031 | 0.520 | 71991 | 0.603 | 1.000 |

**Table S4**: Regression coefficients for the session duration in seconds in the set-shifting experiments, from a linear mixed-effects model with a gamma distribution and identity link function. Bonferroni adjustments are shown applied only to the stimulation coefficients.

Formula: Duration ~ Stimulation:StimulationSite + StimulationSite + Schedule + (1|Subject)

Fixed effects coefficients (95% CIs)

| Name | Coefficient | SE | tstat | DF | p-value | Adjusted.p |
|---|---|---|---|---|---|---|
| (Intercept) | 1307.769 | 105.995 | 12.338 | 540 | 1.000e-26 | |
| Site_dorsolateral | -9.543 | 144.790 | -0.066 | 31 | 0.948 | |
| Site_dorsomedial | 49.190 | 145.074 | 0.339 | 31 | 0.737 | |
| Site_mid | 72.672 | 148.940 | 0.488 | 31 | 0.629 | |
| Schedule_2 | 26.666 | 42.037 | 0.634 | 540 | 0.526 | |
| Schedule_3 | 21.608 | 42.128 | 0.513 | 540 | 0.608 | |
| Schedule_4 | -44.932 | 40.967 | -1.097 | 540 | 0.273 | |
| Stim_ON:Site_ventral | 19.708 | 57.189 | 0.345 | 540 | 0.731 | 1.000 |
| Stim_ON:Site_dorsolateral | 55.030 | 59.901 | 0.919 | 540 | 0.359 | 1.000 |
| Stim_ON:Site_dorsomedial | -14.899 | 58.064 | -0.257 | 540 | 0.798 | 1.000 |
| Stim_ON:Site_mid | 54.967 | 60.899 | 0.903 | 540 | 0.367 | 1.000 |

**Table S5**: Regression coefficients (log scale) for the total number of incorrect responses in the set-shifting experiments, from a generalized linear mixed-effects model with a Negative Binomial distribution and log link function. Bonferroni adjustments are shown applied only to the stimulation coefficients

Formula: Errors ~ Stimulation:StimulationSite + StimulationSite + Schedule + (1|Subject)

Fixed effects coefficients (95% CIs)

| Name | Coefficient | SE | tstat | DF | p-value | Adjusted.p |
|---|---|---|---|---|---|---|
| (Intercept) | 3.735 | 0.058 | 64.772 | 540 | 1.000e-26 | |
| Site_dorsolateral | -0.111 | 0.075 | -1.491 | 31 | 0.146 | |
| Site_dorsomedial | 0.027 | 0.075 | 0.357 | 31 | 0.723 | |
| Site_mid | 0.153 | 0.076 | 2.003 | 31 | 0.054 | |
| Schedule_2 | 0.019 | 0.036 | 0.522 | 540 | 0.602 | |
| Schedule_3 | 0.043 | 0.036 | 1.202 | 540 | 0.230 | |
| Schedule_4 | -0.016 | 0.035 | -0.449 | 540 | 0.653 | |
| Stim_ON:Site_ventral | 0.020 | 0.048 | 0.407 | 540 | 0.684 | 1.000 |
| Stim_ON:Site_dorsolateral | 0.041 | 0.050 | 0.818 | 540 | 0.414 | 1.000 |
| Stim_ON:Site_dorsomedial | -0.066 | 0.049 | -1.340 | 540 | 0.181 | 0.723 |
| Stim_ON:Site_mid | 0.007 | 0.052 | 0.144 | 540 | 0.886 | 1.000 |

**Table S6:**

Regression coefficients for reaction time (RT) as a function of the time (measured in seconds) the rat spent immobile (Immobility) during an open field assay preceding the task from a generalized linear mixed effects model with a gamma distribution and identity link function. Coefficients quantify the effect on RT (measured in seconds).

Formula: RT ~ Immobility + (1|Subject)

Fixed effects coefficients (95% CIs):

| Name | Coefficient | SE | tstat | DF | p-value | Adjusted.p |
|---|---|---|---|---|---|---|
| (Intercept) | 0.63295 | 0.044 | 14.260 | 28386 | 5.599e-46 | |
| Immobility | 4.2873e-05 | 1.38e-05 | 3.107 | 28386 | 1.893e-3 | 7.5728e-3 |

**Table S7:**

Regression coefficients for reaction time (RT) as a function of the distance (measured in cm) the rat traveled (Distance) during an open field assay preceding the task from a generalized linear mixed effects model with a gamma distribution and identity link function. Coefficients quantify the effect on RT (measured in seconds).

Formula: RT ~ Distance + (1|Subject)

Fixed effects coefficients (95% CIs):

| Name | Coefficient | SE | tstat | DF | p-value | Adjusted.p |
|---|---|---|---|---|---|---|
| (Intercept) | 0.66029 | 0.046 | 14.447 | 28386 | 3.823e-47 | |
| Distance | -7.5581e-06 | 5.805e-06 | -1.302 | 28386 | 0.772 | 1.000 |

**Table S8:**

Regression coefficients for reaction time (RT) as a function of the rat's speed (measured in cm/s) during an open field assay preceding the task from a generalized linear mixed effects model with a gamma distribution and identity link function. Coefficients quantify the effect on RT (measured in seconds).

Formula: RT ~ Speed + (1 | Subject)

Fixed effects coefficients (95% CIs):

| Name | Coefficient | SE | tstat | DF | p-value | Adjusted.p |
|---|---|---|---|---|---|---|
| (Intercept) | 0.651 | 0.046 | 14.141 | 28386 | 3.035e-45 | |
| Speed | -5.979e-3 | 0.018 | -0.336 | 28386 | 0.737 | 1.000 |

**Table S9:**

Regression coefficients for reaction time (RT) as a function of a rat's delay in initiating trials (DelayTime, measured in s) from a generalized linear mixed effects model with a gamma distribution and identity link function. Coefficients quantify the effect on RT (measured in seconds).

Formula: RT ~ DelayTime + (1 | Subject)

Fixed effects coefficients (95% CIs):

| Name | Coefficient | SE | tstat | DF | p-value | Adjusted.p |
|---|---|---|---|---|---|---|
| (Intercept) | 0.588 | 0.033 | 17.736 | 26613 | 5.549e-70 | |
| DelayTime | 0.017 | 8.892e-4 | 19.457 | 26613 | 9.696e-84 | 3.879e-83 |

**Table S10:** Regression coefficients for the locomotor measures in the Open Field test, including immobility (seconds), distance traveled (meters), and mean speed (meters/second) and delay time (seconds) – for animals implanted at the mid-striatal site. A Bonferroni correction is shown for the stimulation coefficients as described in the Methods. The coefficients for both the duration of immobility and delay time are derived from a generalized linear mixed-effects model with a gamma distribution and identity link function. The coefficients for both the distance traveled and mean speed are presented on a log scale, originating from log-transformed values to fit a Lognormal distribution, analyzed using a linear mixed-effects model.

General formula: Variable ~ Stimulation + (1|Subject)

Fixed effects coefficients (95% CIs)

| Outcome | Name | Coefficient | SE | tstat | DF | p-value | Adjusted.p |
|---|---|---|---|---|---|---|---|
| Immobility | (Intercept) | 7.446 | 0.072 | 103.378 | 197 | 1.378e-173 | |
| | Stim_ON | -0.019 | 0.020 | -0.924 | 197 | 0.357 | 1.000 |
| Distance | (Intercept) | 303.988 | 28.965 | 10.495 | 197 | 9.440e-21 | |
| | Stim_ON | 23.518 | 14.552 | 1.616 | 197 | 0.108 | 0.431 |
| Speed | (Intercept) | -0.455 | 0.070 | -6.509 | 197 | 6.116e-10 | |
| | Stim_ON | -0.007 | 0.016 | -0.420 | 197 | 0.675 | 1.000 |
| DelayTime | (Intercept) | 1.6781 | 0.14458 | 11.607 | 26613 | 4.5147e-31 | |
| | Stim_ON | -0.0081527 | 0.034242 | -0.23809 | 26613 | 0.81181 | 1.000 |

**Table S11**: Regression coefficients (log scale) for the number of incorrect responses in set-shifting low frequency/high frequency experiments, from a generalized linear mixed-effects model with a Negative Binomial distribution and log link function. Bonferroni adjustments are shown applied only to the stimulation coefficients. Higher coefficients represent more errors, i.e. 20 Hz stimulation caused a significant worsening.

Formula: Errors ~ Stimulation:StimulationSite + StimulationSite + Schedule + (1|StimulationFrequencyGroup:Subject)

Fixed effects coefficients (95% CIs)

| Name | Coefficient | SE | tstat | DF | p-value | Adjusted.p |
|---|---|---|---|---|---|---|
| (Intercept) | 3.595 | 0.094 | 38.222 | 99 | 0 | |
| Stim_frequency_20 | 0.234 | 0.061 | 3.813 | 99 | 1.375e-04 | 5.498e-04 |
| Stim_frequency_130 | -0.003 | 0.033 | -0.100 | 99 | 0.920 | 1.000 |
| Schedule_2 | 0.235 | 0.043 | 5.492 | 99 | 3.980e-08 | |
| Schedule_3 | 0.129 | 0.038 | 3.349 | 99 | 8.116e-04 | |
| Schedule_4 | 0.028 | 0.043 | 0.658 | 99 | 0.511 | |

**Table S12:** Regression coefficients for reaction time (RT) in seconds in the set-shifting low frequency/high frequency experiments, from a generalized linear mixed-effects model, with a gamma distribution and identity link function. Bonferroni adjustments are shown applied only to the stimulation coefficients. In this and subsequent analyses on the same cohort, different Side/Light block sequences (Schedules) had significant behavioral effects. We attribute this to the relatively low N (n=5 rats) in this cohort compared to the main analysis, where the larger cohort (n=36) washed out these effects of no interest. Here, they are regressed out to reveal the primary effects of stimulation.

Formula: RT~ StimulationFrequency + Rule + Schedule + (1|StimulationFrequencyGroup:Subject)

Fixed effects coefficients (95% CIs)

| Name | Coefficient | SE | tstat | DF | p-value | Adjusted.p |
|---|---|---|---|---|---|---|
| (Intercept) | 0.577 | 0.062 | 9.245 | 14855 | 2.352e-20 | |
| Stim_frequency_20 | 0.025 | 0.006 | 3.963 | 14855 | 7.397e-05 | 2.959e-04 |
| Stim_frequency_130 | -0.048 | 0.004 | -12.369 | 14855 | 3.842e-35 | 1.537e-34 |
| Rule_Side | -0.007 | 0.003 | -2.180 | 14855 | 0.029 | |
| Schedule_2 | 0.041 | 0.005 | 8.181 | 14855 | 2.820e-16 | |
| Schedule_3 | 0.018 | 0.004 | 4.238 | 14855 | 2.253e-05 | |
| Schedule_4 | 0.029 | 0.005 | 6.090 | 14855 | 1.130e-09 | |

**Table S13:** Regression coefficients for task accuracy (probability of correct responses) in the set-shifting low frequency/high frequency experiments, from a generalized linear mixed-effects model, with a binomial distribution and logit link function. Bonferroni adjustments are shown applied only to the stimulation coefficients

Formula: Accuracy ~ StimulationFrequency + Rule + Schedule + (1|StimulationFrequencyGroup:Subject)

Fixed effects coefficients (95% CIs)

| Name | Coefficient | SE | tstat | DF | p-value | Adjusted.p |
|---|---|---|---|---|---|---|
| (Intercept) | 1.037 | 0.061 | 16.894 | 14855 | 4.945e-64 | |
| Stim_frequency_20 | -0.036 | 0.079 | -0.451 | 14855 | 0.652 | 1.000 |
| Stim_frequency_130 | -0.032 | 0.042 | -0.765 | 14855 | 0.444 | 1.000 |
| Rule_Side | -0.580 | 0.036 | -15.929 | 14855 | 4.007e-57 | |
| Schedule_2 | -0.120 | 0.054 | -2.207 | 14855 | 0.027 | |
| Schedule_3 | -0.054 | 0.048 | -1.124 | 14855 | 0.261 | |
| Schedule_4 | 0.084 | 0.054 | 1.548 | 14855 | 0.122 | |

**Table S14:** Regression coefficients for the session duration in seconds in set-shifting low frequency/high frequency experiments, from a linear mixed-effects model. Bonferroni adjustments are shown applied only to the stimulation coefficients. A longer duration implies worse performance, i.e. we see a significant worsening induced by 20 Hz stimulation.

Formula: Duration ~ StimulationFrequency + Schedule + (1|StimulationFrequencyGroup:Subject)

Fixed effects coefficients (95% CIs)

| Name | Coefficient | SE | tstat | DF | p-value | Adjusted.p |
|---|---|---|---|---|---|---|
| (Intercept) | 1090.760 | 75.695 | 14.410 | 99 | 4.477e-47 | |
| Stim_frequency_20 | 188.395 | 65.899 | 2.859 | 99 | 0.004 | 0.017 |
| Stim_frequency_130 | -56.208 | 57.985 | -0.969 | 99 | 0.332 | 1.000 |
| Schedule_2 | 188.266 | 61.056 | 3.084 | 99 | 0.002 | |
| Schedule_3 | 118.433 | 54.622 | 2.168 | 99 | 0.030 | |
| Schedule_4 | 106.741 | 59.434 | 1.796 | 99 | 0.073 | |

**Table S15:** Training schedule for the 5-choice serial reaction time task, progressively increasing in difficulty. Training stage denotes difficulty level, stimulus duration indicates how long the correct stimulus was shown to the rat, inter-trial interval indicates the time between initiation and when the stimulus is shown, limited hold indicates the time the rat had to make a decision after the stimulus was shown, and criterion indicates the conditions the rat must meet to continue to the next training stage.

| Training Stage | Stimulus Duration (s) | Initiation to Stimulus Delay (s) | Limited Hold (s) | Criterion |
|---|---|---|---|---|
| 1 | 30 | 2 | 30 | ≥ 30 Correct trials |
| 2 | 20 | 2 | 20 | ≥ 30 Correct trials |
| 3 | 10 | 5 | 10 | ≥ 50 Correct trials<br>≥ 80% Accuracy<br>≥ 20% Omissions |
| 4 | 5 | 5 | 5 | ≥ 50 Correct trials<br>≥ 80% Accuracy<br>≥ 20% Omissions |
| 5 | 2.5 | 5 | 5 | ≥ 50 Correct trials<br>≥ 80% Accuracy<br>≥ 20% Omissions |
| 6 | 1.25 | 5 | 5 | ≥ 50 Correct trials<br>≥ 80% Accuracy<br>≥ 20% Omissions |
| 7 | 1 | 5 | 5 | ≥ 50 Correct trials<br>≥ 80% Accuracy<br>≥ 20% Omissions |
| 8 | 0.9 | 5 | 5 | ≥ 50 Correct trials<br>≥ 80% Accuracy |

| | | | | |
|---|---|---|---|---|
| | | | | ≥ 20% Omissions |
| 9 | 0.8 | 5 | 5 | ≥ 50 Correct trials<br>≥ 80% Accuracy<br>≥ 20% Omissions |
| 10 | 0.7 | 5 | 5 | ≥ 50 Correct trials<br>≥ 80% Accuracy<br>≥ 20% Omissions |
| 11 | 0.6 | 5 | 5 | ≥ 50 Correct trials<br>≥ 80% Accuracy<br>≥ 20% Omissions |
| 12 | 0.5 | 5 | 5 | ≥ 50 Correct trials<br>≥ 80% Accuracy<br>≥ 20% Omissions |

**Table S16:** Number of sessions required for rats to pass all training stages for the 5-choice serial reaction time task. The first number indicates the number of sessions to pass for the first time while the second indicates the number of sessions needed to pass once again after surgery.

| Animal | # of sessions to pass Training Stage 12 |
|---|---|
| FC2 | 64+10 |
| FC3 | 49+11 |
| FC30 | 167+24 |
| FC31 | 257+3 |
| FC32 | 99+9 |
| FC35 | 90+24 |

**Table S17:** Regression coefficients for the probability of premature responses in the 5-choice serial reaction time task, from a generalized linear mixed-effects model, with a binomial distribution and logit link function. Bonferroni adjustment is shown applied only to the stimulation coefficient. This and subsequent 5CSRTT analyses only describe mid-striatal stimulation.

Formula: Premature ~ Stim + Schedule + (1|Subject)

Fixed effects coefficients (95% CIs)

| Name | Coefficient | SE | tstat | DF | p-value | Adjusted.p |
|---|---|---|---|---|---|---|
| (Intercept) | -1.730 | 0.135 | -12.810 | 8218 | 3.283e-37 | |
| Stim_ON | 0.106 | 0.062 | 1.716 | 8218 | 0.086 | 0.345 |
| Schedule_2 | 0.005 | 0.091 | 0.055 | 8218 | 0.956 | |
| Schedule_3 | 0.028 | 0.092 | 0.307 | 8218 | 0.759 | |
| Schedule_4 | -0.163 | 0.095 | -1.718 | 8218 | 0.086 | |
| Schedule_5 | -0.018 | 0.092 | -0.193 | 8218 | 0.847 | |

**Table S18:** Regression coefficients for RT in the 5-choice serial reaction time task, from a generalized linear mixed-effects model, with a gamma distribution and identity link function. Bonferroni adjustment is shown applied only to the stimulation coefficient. A positive coefficient for stimulation indicates a slowing of RT, in contrast to the speeding seen during Set Shift with mid-striatal stimulation.

Formula: RT ~ Stim + Schedule + (1|Subject)

Fixed effects coefficients (95% CIs)

| Name | Coefficient | SE | tstat | DF | p-value | Adjusted.p |
|---|---|---|---|---|---|---|
| (Intercept) | 0.618 | 0.041 | 14.908 | 6867 | 1.738e-49 | |
| Stim_ON | 0.049 | 0.015 | 3.212 | 6867 | 0.001 | 0.005 |
| Schedule_2 | 0.041 | 0.024 | 1.751 | 6867 | 0.080 | |
| Schedule_3 | -0.012 | 0.022 | -0.517 | 6867 | 0.605 | |
| Schedule_4 | -0.018 | 0.022 | -0.831 | 6867 | 0.406 | |
| Schedule_5 | -0.031 | 0.022 | -1.413 | 6867 | 0.158 | |

**Table S19:** Regression coefficients for the probability of omissions in the 5-choice serial reaction time task, from a generalized linear mixed-effects model, with a binomial distribution and logit link function. Bonferroni adjustment is shown applied only to the stimulation coefficient.

Formula: Omission ~ Stim + Schedule + (1|Subject)

Fixed effects coefficients (95% CIs)

| Name | Coefficient | SE | tstat | DF | p-value | Adjusted.p |
|---|---|---|---|---|---|---|
| (Intercept) | -2.689 | 0.219 | -12.286 | 9068 | 2.021e-34 | |
| Stim_ON | 0.448 | 0.075 | 5.958 | 9068 | 2.653e-09 | 1.061e-08 |
| Schedule_2 | -0.041 | 0.115 | -0.359 | 9068 | 0.720 | |
| Schedule_3 | 0.249 | 0.110 | 2.261 | 9068 | 0.024 | |
| Schedule_4 | 0.015 | 0.113 | 0.132 | 9068 | 0.895 | |
| Schedule_5 | 0.101 | 0.113 | 0.897 | 9068 | 0.370 | |

**Table S20:** Regression coefficients for accuracy in the 5-choice serial reaction time task, from a generalized linear mixed-effects model, with a binomial distribution and logit link function. Bonferroni adjustment is shown applied only to the stimulation coefficient

Formula: Accuracy ~ Stim + Schedule + (1|Subject)

Fixed effects coefficients (95% CIs)

| Name | Coefficient | SE | tstat | DF | p-value | Adjusted.p |
|---|---|---|---|---|---|---|
| (Intercept) | 1.405 | 0.207 | 6.780 | 6918 | 1.297e-11 | |
| Stim_ON | -0.007 | 0.060 | -0.123 | 6918 | 0.902 | 1.000 |
| Schedule_2 | -0.121 | 0.089 | -1.355 | 6918 | 0.175 | |
| Schedule_3 | 0.003 | 0.092 | 0.035 | 6918 | 0.972 | |
| Schedule_4 | 0.077 | 0.092 | 0.838 | 6918 | 0.402 | |
| Schedule_5 | -0.033 | 0.090 | -0.364 | 6918 | 0.716 | |

**Table S21:** Regression coefficients for the number of pokes into the previous trial's correct port made during the ITI (correctEarly/Late), as a function of whether the rat made a correct choice in the preceding trial (accuracy), from a generalized linear mixed-effects models with a Poisson distribution and identity link function. Positive accuracy coefficients represent repetition of a correct response, whereas negative coefficients represent corrective behavior (a tendency to poke the port only when it was not chosen correctly during the preceding trial).

Formula: correctEarly/Late ~ accuracy + (1 | Subject)

Fixed effects coefficients (95% CIs):

| Outcome | Name | Coefficient | SE | tstat | DF | p-value |
|---|---|---|---|---|---|---|
| correctEarly | (Intercept) | 0.267 | 0.005 | 52.934 | 28064 | 0 |
| | accuracy | -0.238 | 0.005 | -45.783 | 28064 | 0 |
| correctLate | (Intercept) | 0.287 | 0.005 | 54.918 | 3803 | 0 |
| | accuracy | 0.115 | 0.007 | 16.202 | 3803 | 9.054e-59 |

**Table S22:** Regression coefficients for the proportion of left pokes made during the early or late ITI, as a function of whether the left port was the correct option on the preceding trial (leftCorrect), from a generalized linear mixed-effects models with a binomial distribution and logit link function. This model was restricted to ITIs following incorrect responses, i.e. positive coefficients for leftCorrect would signify corrective behavior (poking the left port because it would have been rewarded).

Formula: leftMostEarly/Late ~ leftCorrect + (1 | Subject)

Fixed effects coefficients (95% CIs):

| Outcome | Name | Coefficient | SE | tstat | DF | p-value |
|---|---|---|---|---|---|---|
| leftMostEarly | (Intercept) | -0.185 | 0.081 | -2.300 | 4535 | 0.021 |
| | leftCorrect | 0.527 | 0.061 | 8.649 | 4535 | 7.094e-18 |
| leftMostLate | (Intercept) | -0.7716 | 0.136 | -5.673 | 4002 | 1.505e-8 |
| | leftCorrect | 1.726 | 0.071 | 24.218 | 4002 | 4.940e-121 |

**Table S23:** Regression coefficients for the proportion of left pokes made during the early or late ITI, as a function of whether the left port was the correct option on the preceding trial (leftCorrect), from a generalized linear mixed-effects models with a binomial distribution and logit link function. This model was restricted to ITIs following correct responses, i.e. it cannot show corrective behaviors. Positive coefficients signify repetition of a correct choice.

Formula: leftMostEarly/Late ~ leftCorrect + (1 | Subject)

Fixed effects coefficients (95% CIs):

| Outcome | Name | Coefficient | SE | tstat | DF | p-value |
|---|---|---|---|---|---|---|
| leftMostEarly | (Intercept) | -4.206 | 0.614 | -6.846 | 687 | 1.690e-11 |
| | leftCorrect | 8.269 | 0.651 | 12.700 | 687 | 2.407e-33 |
| leftMostLate | (Intercept) | -0.926 | 0.137 | -6.779 | 8814 | 1.290e-11 |
| | leftCorrect | 1.886 | 0.049 | 38.636 | 8814 | 7.982e-302 |

**Table S24:** Regression coefficients for choosing the left port (leftChoice) as a function of whether the left port was poked most frequently during the end of the preceding ITI (moreLeftLate), from a generalized linear mixed-effects models with a binomial distribution and logit link function. Separate models were run for trials where the illuminated port was and was not chosen, because we expected rats to commit to different strategies on these trials. The relatively small negative coefficient for the Light Chosen model implies that ITI rehearsals do not strongly shape choices during behavior when the rat is focused on the cue light. (This may be during a Light rule, or during a Side rule when the rat is erroneously using light-focused behavior.) Instead, rats were slightly less likely to poke a rehearsed port. In contrast, a much larger moreLeftLate coefficient for the Light Not Chosen model highlights that outside of light rule behavior, late ITI pokes have a strong effect on the subsequent trial's choice.

Formula: leftChoice ~ moreLeftLate + (1 | Subject)

Fixed effects coefficients (95% CIs):

| Group | Name | Coefficient | SE | tstat | DF | p-value |
|---|---|---|---|---|---|---|
| Light Chosen | (Intercept) | 0.252 | 0.031 | 8.143 | 9076 | 4.373e-16 |
| | moreLeftLate | -0.087 | 0.042 | -2.051 | 9076 | 0.080 |
| Light Not Chosen | (Intercept) | -0.852 | 0.085 | -10.040 | 3803 | 1.992e-23 |
| | moreLeftLate | 1.844 | 0.073 | 25.131 | 3803 | 4.499e-129 |

**Table S25:** Regression coefficients for reaction time (RT) as a function of behaviors during the end of the preceding ITI, from a generalized linear mixed effects model with a gamma distribution and identity link function. Coefficients quantify the effect on RT (measured in seconds) of a poke into the middle (trial initiation) port vs. that poke combined with a side port poke. nonchoiceMostLate captures pokes into a port that will not be chosen on the upcoming trial, while choiceMostLate captures pokes into the port that will be chosen. These are mutually exclusive indicators, signifying which side port received more pokes during the final 2 seconds of the ITI. Negative coefficients in any of these terms imply that that behavior produces faster RTs. Thus, the results show that middle port pokes and rehearsal pokes (choiceMostLate) both speed RT.

Formula: RT ~ midPokeLate + nonchoiceMostLate:midPokeLate + choiceMostLate:midPokeLate + (1 | Subject)

Fixed effects coefficients (95% CIs):

| Name | Coefficient | SE | tstat | DF | p-value |
|---|---|---|---|---|---|
| (Intercept) | 0.665 | 0.044 | 15.045 | 28446 | 5.892e-51 |
| midPokeLate | -0.036 | 4.546e-3 | -7.969 | 28446 | 1.656e-15 |
| nonchoiceMostLate: midPokeLate | 4.458e-3 | 5.834e-3 | -0.764 | 28446 | 0.445 |
| choiceMostLate: midPokeLate | -0.012 | 5.251e-3 | -2.306 | 28446 | 0.021 |

**Table S26:** Regression coefficients for the probability of having at least one of various types of poke in the late portion of the ITI as a function of active/sham mid-striatal stimulation, from a generalized linear mixed effects model with a binomial distribution and logit link function. A significant positive coefficient indicates that stimulation increases the probability of that behavior. Given that stimulation increases all types of ITI pokes, it cannot be characterized as specifically affecting proactive cognitive control or rehearsal. Those would lead to increases in Choice pokes but not Non-choice pokes.

Formula: PokeType ~ Stim + (1|Subject)

Fixed effects coefficients (95% CIs):

| Poke Type | Name | Coefficient | SE | tstat | DF | p-value |
|---|---|---|---|---|---|---|
| Mid | (Intercept) | -0.474 | 0.154 | -3.076 | 28448 | 2.075e-3 |
| | Stim | 0.237 | 0.025 | 9.487 | 28448 | 2.556e-21 |
| Choice | (Intercept) | -1.219 | 0.109 | -11.164 | 28448 | 7.049e-29 |
| | Stim | 0.206 | 0.028 | 7.481 | 28448 | 7.619e-14 |
| Non-choice | (Intercept) | -1.655 | 0.104 | -15.898 | 28448 | 1.495e-56 |
| | Stim | 0.087 | 0.032 | 2.726 | 28448 | 6.415e-3 |
